# Bacterial targeting of host paraspeckles uncovers a new SFPQ-based regulation

**DOI:** 10.64898/2026.01.11.698932

**Authors:** Sonia Nicchi, Sonia Mondino, Christophe Rusniok, Nassim Mahtal, Bertrand Raynal, Rayen Elj, Quentin Giai Gianetto, Joaquín Dalla Rizza, Nicole Larrieux, Jessica E. Martyn, Hanno Schoeler, Sandrine Schmutz, Florian Muller, Mariette Matondo, Alejandro Buschiazzo, Carmen Buchrieser, Monica Rolando

## Abstract

Nuclear functions are key in protecting cells against infections, yet intracellular pathogens like *Legionella pneumophila* can exploit these mechanisms to survive. We characterized a *L. pneumophila* protein, LpDot1, which shares sequence similarity with the eukaryotic catalytic domain of the histone methyltransferase DOT1. Structure determination, together with biochemical and biophysical analyses, revealed that LpDot1 methylates non-histone nuclear proteins, notably the splicing factor proline-glutamine rich protein (SFPQ). Importantly, LpDot1 targets the previously uncharacterized K518, located on an important structural motif of SFPQ, therefore impairing its dimerization *in vitro*. During infection, *L. pneumophila* modulates SFPQ abundance and activities in a LpDot1-dependent manner, thereby hijacking paraspeckle organization and the host cell splicing machinery, leading to alternative splice variants of infection related genes such as NF-kB2 and CD45. To our knowledge, this is the first report of a bacterial effector directly modifying paraspeckle dynamics, providing new insight into previously uncharacterized eukaryotic regulatory pathways.

## INTRODUCTION

The nucleus is a highly organized and dynamic organelle that controls various cellular activities, including gene expression, RNA processing, and DNA replication^1^. The term ’nucleus,’ derived from the Latin ’kernel’ or ’seed,’ reflects its role as command center of the cell^2^. It is enclosed by the nuclear envelope, which contains nuclear pore complexes that regulate transport into and out of the nucleus^3^. Chromatin, located within the nucleus, is a nucleoprotein complex consisting of DNA wrapped around histone octamers. The nucleoplasm is populated by various membraneless organelles (MLOs), including nucleoli, speckles, and paraspeckles. These play essential roles in RNA metabolism, gene regulation, and stress responses ^4^. First described in HeLa cells in 2002, paraspeckles are ribonucleoprotein bodies of 0.5-2.0 µm in size that are found in the interchromatin space close to but distinct from the nuclear speckles^5^. They are composed of a shell and a core where the lncRNA NEAT1 (nuclear-enriched autosomal transcript-1)^6^ is critical for paraspeckle formation and maintenance, facilitating the assembly of stable ribonucleoprotein particles. Paraspeckles contain a small number of well identified RNA-binding proteins belonging to the Drosophila Behavior Human Splicing (DBHS) family proteins, such as FUS, PSPC1, NONO, and the splicing factor proline-glutamine rich protein (SFPQ)^7^ ^8^ ^9^. SFPQ is a multifunctional protein involved in paraspeckle formation, RNA biogenesis, transport, and the regulation of alternative splicing^10^, a complex process that generates multiple mRNA variants from a single gene. Paraspeckles are highly dynamic structures that vary in number, size and morphology in response to cellular stress conditions. Importantly, dysregulation of paraspeckles and aberrant splicing have been reported in numerous diseases, including cancer and viral infections^11^ ^12^, underscoring the importance of these structures in maintaining cellular homeostasis^13^. Recent findings also highlighted that the dynamic nature of paraspeckles plays a crucial role in inflammatory gene expression following stress stimuli such as bacterial lipopolysaccharide (LPS)^14^.

Although it has never been described that bacterial pathogens target paraspeckles, it is well known that they can target the nucleus through effector proteins promoting replication and persistence within infected host cells^15^. These nuclear-targeted bacterial proteins have been designated “nucleomodulins” ^16^. *Legionella pneumophila,* a gram-negative bacterial pathogen causing community- and hospital acquired pneumonia called Legionnaire’s disease is one of these bacteria that targets the nucleus via its Dot/Icm type-4 (T4SS) secreted effector proteins^17^. Most interestingly, during the long-lasting coevolution with protozoa, *Legionella* spp. have evolved survival mechanisms many of which are conferred by a vast repertoire of eukaryotic-like proteins acquired through horizontal gene transfer from their protozoan hosts ^18,19^, some of which encode eukaryotic, nuclear functions ^20,21^.

Here, we functionally characterized a novel *L. pneumophila* effector protein, that encodes lysine methyltransferase activity. We named it LpDot1 due to its homology with the catalytic domain of the eukaryotic DOT1 (Disruptor of Telomeric Silencing-1) reported to exclusively methylate lysine K79 in the globular region of histone H3, affecting gene expression and DNA damage response^22^. Unexpectedly, LpDot1 does not methylate histones but functions within macrophages and methylates nuclear proteins, thereby altering paraspeckle composition and alternative splicing of the human host cell. This posttranslational modification discovered here, enhances intracellular replication of *L. pneumophila* through modulation of alternative splicing thereby dampening the host immune response.

## RESULTS

### *Legionella pneumophila* encodes a eukaryotic Dot1-like protein

Genome analyses of *L. pneumophila* revealed a gene (*lpp3025*) predicted to encode a lysine methyltransferase (MTase) similar to the catalytic domain of the eukaryotic DOT1 (Disruptor of Telomeric Silencing-1). Importantly, this gene is conserved in the genus *Legionella* as it is present among all 58 different *Legionella* species sequenced, suggesting it encodes an important function^19^. Given its high similarity to DOT1 we named the *Legionella* protein LpDot1. It contains the catalytic signature of DOT1 enzymes (black boxes in **Fig 1A** and **Fig. S1**), notably the DxGxGxG motif necessary for SAM (S-adenosylmethionine) binding and essential for the catalytic activity^23^. Local Pairwise Sequence Alignment (LALIGN) helped to evaluate the percentage of aminoacidic identity and similarity between the LpDot1 protein and its human homolog DOT1L, as well as with selected eukaryotic homologs present in the genomes of the environmental hosts of *L. pneumophila*, such as *Acanthamoeba castellanii*, and other protists present in the P10K database^24^. As illustrated in **Fig. 1A**, the percentages of aminoacidic identity and similarity between LpDot1 and its eukaryotic homologs is high but variable, ranging from 55.9 - to 77.7% of similarity and from 22.9- to 48.7% of identity. Interestingly, among the top three matches, we found two ciliates (*Telotrochidium* spp. and *Sessilida* spp.) and one alga (*Astrophomene gubernaculifera*) with the high aminoacidic identity of 43.3% and a similarity over 70 %. To note, the *A. castellanii* genome encodes two DOT1-like proteins: a long isoform that closely resembles human DOT1L, as it contains several auxiliary domains in addition to the catalytic domain, and a shorter isoform that consists only of the catalytic methyltransferase domain, mirroring the structure seen in *Legionella* spp. (**Fig 1A**). These analyses strongly substantiate our phylogenetic analyses that the *dot1* gene was acquire via interdomain horizontal gene transfer from a protozoan host sharing the same ecological niche^25^.

**Figure 1:**
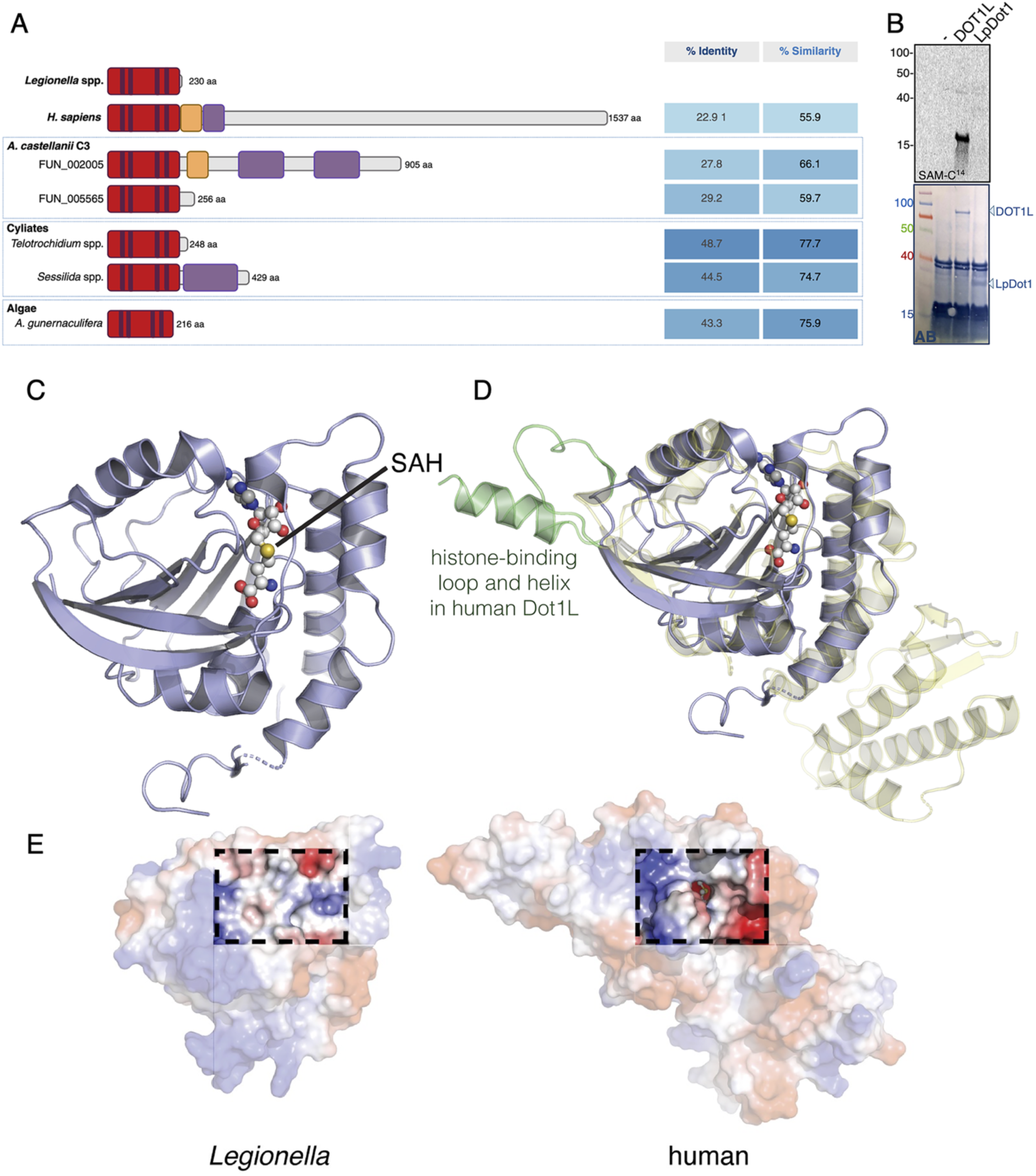
Legionella *pneumophila* encodes a eukaryotic Dot1-like methyltransferase with structural divergences from human DOT1L. (**A**) Schematic representation of *L. pneumophila* LpDot1 and other eukaryotic DOT1-like proteins (*Homo sapiens*[Q8TEK3], *Acanthamoeba castellanii* Strain C3 [FUN_002004, FUN_005565], *Telotrochidium* [P10K-MW-000057|gene13722], *Sessilida* [P10K-MW-000059|gene19854], and *Astrophomene gubernaculifera* [P10K-MW-000389|gene4536]). InterPro and PROSCAN were used to identify conserved motifs or domains (red: DOT1 catalytic domain with dark lines: catalytic conserved motifs; yellow: nuclear localization signal (NLS); purple: variable nuclear binding domains. (**B**) *In vitro* MTase assay of LpDot1 or human DOT1L recombinant proteins with calf thymus histones as substrate in the presence of ^14^C-SAM as a methyl donor. *Upper panel*: ^14^C radiography; *Bottom panel* amido black (AB) staining. (**C**) Crystal structure of LpDot1 (blue cartoon) with bound S-adenosyl-L-homocysteine (SAH, ball-and-stick representation) showing the conserved Class I SAM-dependent methyltransferase fold. **(D)** Structural superposition of LpDot1 with human DOT1L (yellow/green semi-transparent cartoon). Note the absence of critical histone-binding regions in the bacterial enzyme: a 14-residue loop and extended C-terminal α-helix (highlighted in green) essential for nucleosome binding in human DOT1L. **(E)** Electrostatic potential surfaces of bacterial (*left*) and human (*right*) enzymes. Dashed boxes highlight substrate-binding regions, showing highly charged features optimized for histone interactions in human DOT1L (bound SAH in ball-and-stick) versus LpDot1’s neutral profile with distinct gate loops closing the SAM-binding pocket

### LpDot1 exhibits substrate recognition surfaces that are divergent from histone methyltransferases

To determine whether LpDot1 catalyses methyltransferase (MTase) activity similarly to its human homolog, we performed *in vitro* MTase assays with recombinant LpDot1 and calf thymus histones. Differently from the human DOT1L, we observed no detectable MTase activity on histones (**Fig. 1B**). To understand this result and further elucidate its function we determined the three-dimensional structure of LpDot1 (**Table S1**) co-crystallized with SAM. Crystals grew in the trigonal P3₁21 space group and diffracted X-rays to 2.1 Å resolution. LpDot1 adopts an overall architecture similar to the catalytic domain of eukaryotic DOT1 MTases, exhibiting a conserved Class I SAM-dependent MTase fold (**Fig. 1C**). Despite being co-crystallized with SAM, S-adenosyl-L-homocysteine (SAH) was bound within the active site (**Fig. S2A**), suggesting that methyl transfer reactions occurred during crystallization, as reported for other SAM-dependent MTases^26^. SAH occupies a deep binding pocket, featuring the highly conserved Motif I (residues 102-108), in contact with the ribose and methionyl moieties of the ligand. In solution LpDot1 behaves as a monomer, consistent with the crystallographic arrangement (**Fig. S2B-2E**). Comparative analyses with known eukaryotic DOT1 structures (PDBs 1U2Z, 1NW3 and 6J99) revealed significant differences in the substrate-binding regions, strongly suggesting that LpDot1 has adapted to methylate targets other than histones. Two critical structural elements of human DOT1L to bind and methylate histones are absent in LpDot1: (*i*) a loop of 14 amino acids, spanning residues 273-286; and (*ii*) an extended C-terminal α-helix corresponding to residues 319-331 (**Fig. 1D**). These elements provide critical interaction surfaces with human histones H2A and H2B within the nucleosome^27^, and their absence in LpDot1 would preclude stable nucleosome binding. Additionally, the C-terminal helix in human DOT1L is essential for ubiquitin binding, which significantly enhances enzymatic activity by stabilizing the active conformation and reducing the K_M_ for nucleosomal substrates^28^. While human DOT1L presents a highly charged exposed surface, optimal for nucleosomal histone interactions, LpDot1 exhibits a relatively neutral surface (**Fig. 1E**), which includes two key loops that are well positioned to function as substrate-entering gates: loop-EF (residues 70-79: RDAIEYVYGE) and loop-8/J (residues 176-181: INSSTL). In contrast to the rest of the protein, these two loops were predicted with low reliability indices by AlphaFold2, exhibiting significant shifts with respect to the experimental structure. The AF-predicted apo structure displays the gate loops open, which may shift to the closed configuration observed in the crystal, once the SAM substrate enters the binding pocket. Yet another important difference between the human and bacterial proteins is the presence of A72 in LpDot1 gate Loop-EF, replacing F131 in human DOT1L (**Fig 1D**). In DOT1L, F131 works together with W305 and the H4 tail to stabilize the active conformation of histone H3, promoting correct orientation of H3K79 for catalysis^29^ and its substitution by alanine precludes H3-methylation activity in the human enzyme^29^. Taken together, the first experimentally determined 3D structure of a DOT-like MTase of bacterial origin strongly suggests that *L. pneumophila* repurposed a Dot1-like methyltransferase to target host proteins other than histones.

### LpDot1 is an active methyltransferase that targets nuclear non-histone proteins

To test the hypothesis of LpDot1 being an active MTase with repurposed activity and further elucidate the identity of its substrate(s), we performed a MTase assay using human nuclear extract (NE). Indeed, LpDot1 possessed strong MTase activity when incubated with human NE (**Fig. 2A**), suggesting that LpDot1 indeed exhibits a redirected substrate specificity.

**Figure 2.**
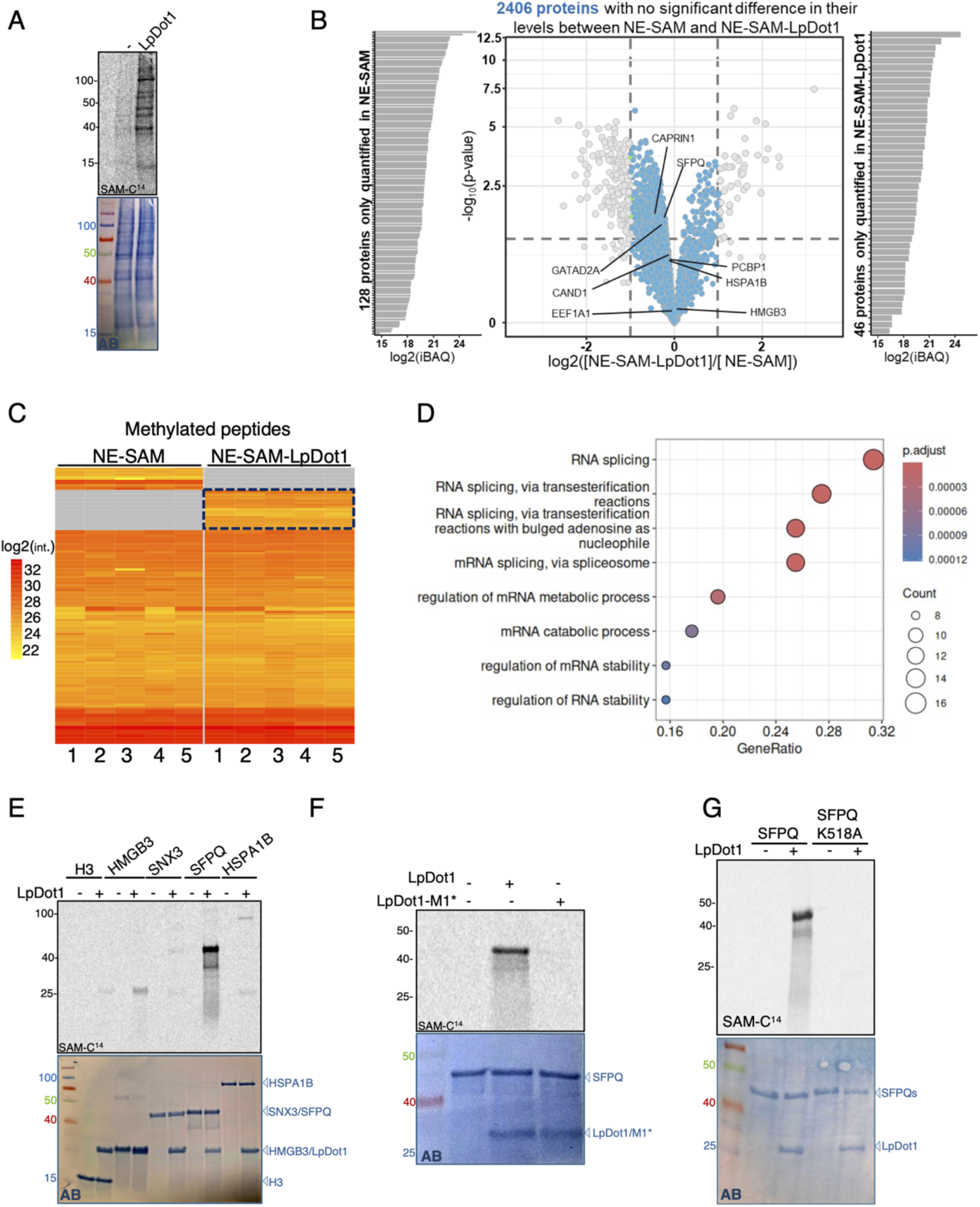
LpDot1 targets nuclear non-histone proteins and uniquely methylates K518 of SFPQ. (**A**) MTase assay of LpDot1 recombinant protein using human nuclear extracts (NE) as substrate in the presence of ^14^C-SAM as a methyl donor. *Upper panel*: ^14^C radiography; *Bottom panel* amido black (AB) staining. (**B**) Liquid chromatography and mass spectrometry (LC-MS) identifying LpDot1 targets. Volcano plot of -log_10_ (p values) vs log_2_ values (NE-SAM-LpDot1/NE-SAM). Blues dots represent proteins which abundance is not different between the two groups of comparison. Gray dots are proteins that are differentially abundant in the two conditions and are represented on vertical bar plots on each side. Results from five independent biological replicates (n=5). (**C**) Heat map showing the differentially methylated peptides in NE-SAM compared to NE-SAM-LpDot1 (n=5). Black box indicates the peptides showing a methylation that is LpDot1-dependent. (**D**) GO analysis of the biological processes enriched in the 64 methylated proteins. (**E**) *In vitro* MTase assay of LpDot1 and the human recombinant proteins: H3 [Q6NXT2], HMGB3 [O15347], SNX3 [O60493], SFPQ [P23246], and HSPA1B [P0DMV9]. (**F**) *In vitro* MTase assay of LpDot1 and LpDot1-M1* with human SFPQ as substrate in the presence of ^14^C-SAM as a methyl donor. (**G**) *In vitro* MTase assay of LpDot1 using either SFPQ or SFPQ-K518A recombinant proteins as substrate in the presence of ^14^C-SAM as a methyl donor. For (**E - G**): *Upper panel*: ^14^C radiography; *Bottom panel* amido black (AB) staining.

To identify the LpDot1 methylation targets, we incubated human NE with or without LpDot1 in the presence of cold SAM, processed samples by liquid chromatography–mass spectrometry (LC-MS), and compared methylation events between the two groups (**Fig. S3A)**. To minimize the impact of protein abundance variations on methylation measurements, and thus ensure accurate quantification of methylation, we analysed 2406 proteins (among the 2580 identified ones) that showed comparable abundance in the two experimental conditions (**Fig 2B**). Sixty-four proteins showed different levels of methylation (**Fig. 2C**), of which 14 were methylated only in the presence of LpDot1 (**Table 1**). Gene Ontology (GO) analysis revealed that these proteins are implicated in RNA splicing *via* transesterification reactions or *via* spliceosome, mRNA metabolic process, and RNA processing (**Fig. 2D**). Interestingly, when the same experimental approach was used with amoeba nuclear extracts as substrate of LpDot1 we found 19 proteins exclusively methylated in the presence of LpDot1(**Fig. S3B** and **Fig. S3C**). Again, mainly RNA-related processes were found to be enriched among the differentially methylated proteins, further supporting the implication of LpDot1 in these biological processes (**Fig. S3D**).

**Table 1.**
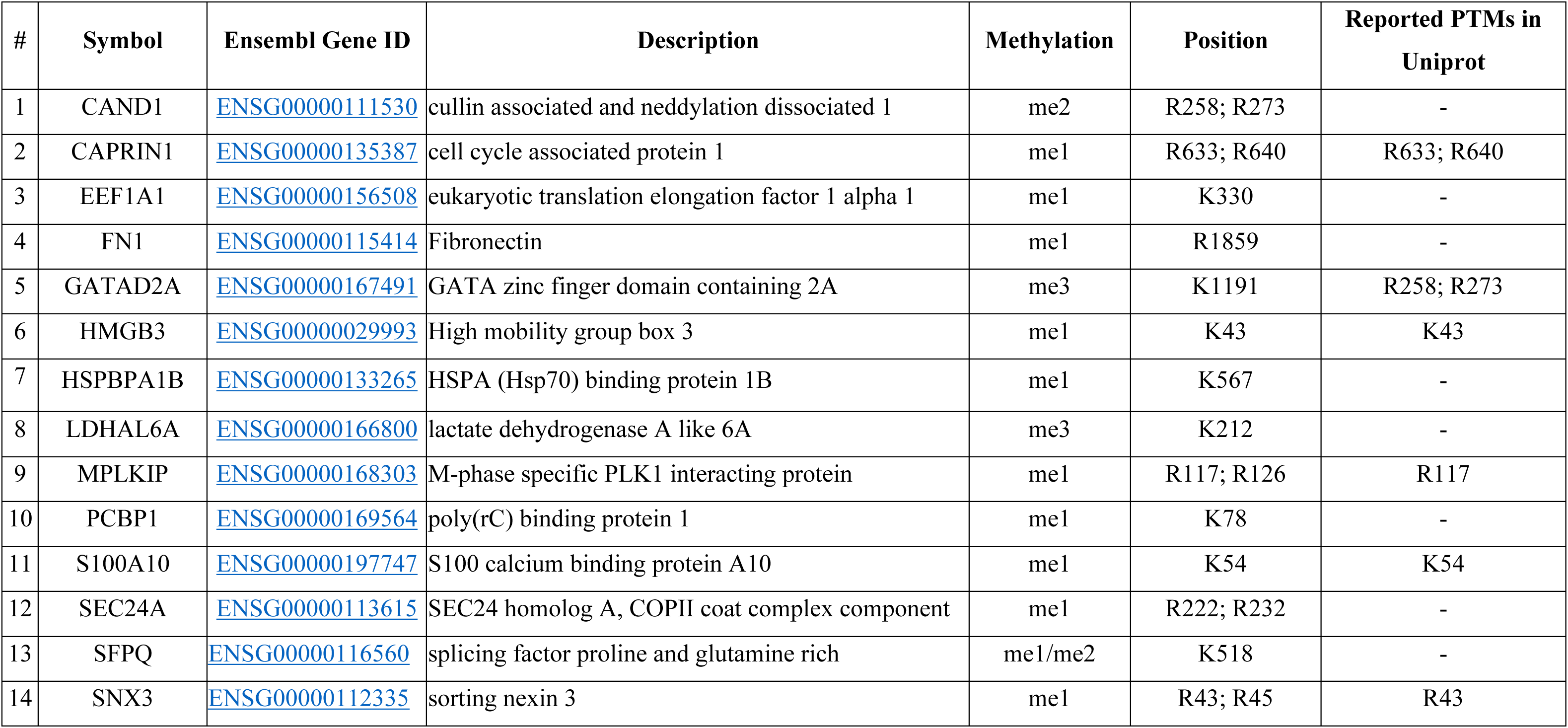
List of the 14 proteins showing a LpDot1-dependent methylation (alphabetic order).

### LpDot1 targets K518 on SFPQ

We chose to experimentally validate a few targets among the proteins identified by MS/MS (**Table 1**). According to their implication in important biological processes and the significant levels of differential methylation we selected the high mobility group protein B3 (HMGB3), the sorting nexin protein (SNX3), the splicing factor proline- and glutamine-rich (SFPQ) and the heat shock protein (HSPA1B). *In vitro* MTase assays confirmed that the four tested, recombinant proteins are methylated by LpDot1, whereby SFPQ showed the strongest methylation profile (**Fig. 2E**). To confirm that LpDot1 specifically catalyzes SFPQ methylation, we mutated the SAM-binding site Motif-I in LpDot1 (shown in **Fig. S1**), a site that has been used to create catalytically inactive variants of human DOT1L^30^. Indeed, mutations at glycine residues 96 and 98 (G96R/G98R) generated a catalytic inactive mutant (LpDot1-M1*) (**Fig. 2F**).

Mass spectrometry analysis of human nuclear extracts incubated with LpDot1 (**Fig. 2B-2D** and **Table 1**) identified K518 of SFPQ, a residue not known previously to be methylated on SFPQ, as the unique target of LpDot1 (**Fig. 2C** and **Table 1**). This site was confirmed by LC-MS using the human recombinant protein as a substrate for LpDot1 (**Fig. S4A**). To further validate this methylation site, we mutated K518 of SFPQ to alanine and repeated the MTase assay with LpDot1. The SFPQ K518A mutant protein was no longer methylated by LpDot1 further confirming that K518 is the unique site methylated by LpDot1(**Fig. 2G**).

### LpDot1 methylation of K518 of SFPQ impairs its dimerization

The structure of SFPQ is characterized by the presence of two low-complexity regions at both the N- and C-termini, intrinsically disordered and rich in proline and glutamine residues that enable SFPQ to act as a scaffold protein by driving phase separation^31^ (**Fig. 3A)**. It also contains different domains responsible for RNA- and DNA-binding, and a conserved region important for the interaction with other members of the DBHS family. A C-terminal coiled-coil region spanning amino acids 528 to 555 possesses an unusual antiparallel configuration that extends over 260 Å that has been reported to be critical for SFPQ oligomerization^31^ ^32^. Intriguingly, K518 is located in this motif, suggesting a possible destabilization of the dimer upon methylation, as other, reported mutations in this motif resulted in SFPQ mislocalization, reduced formation of nuclear bodies, abrogated molecular interactions and deficient transcriptional regulation^31^.

**Figure 3.**
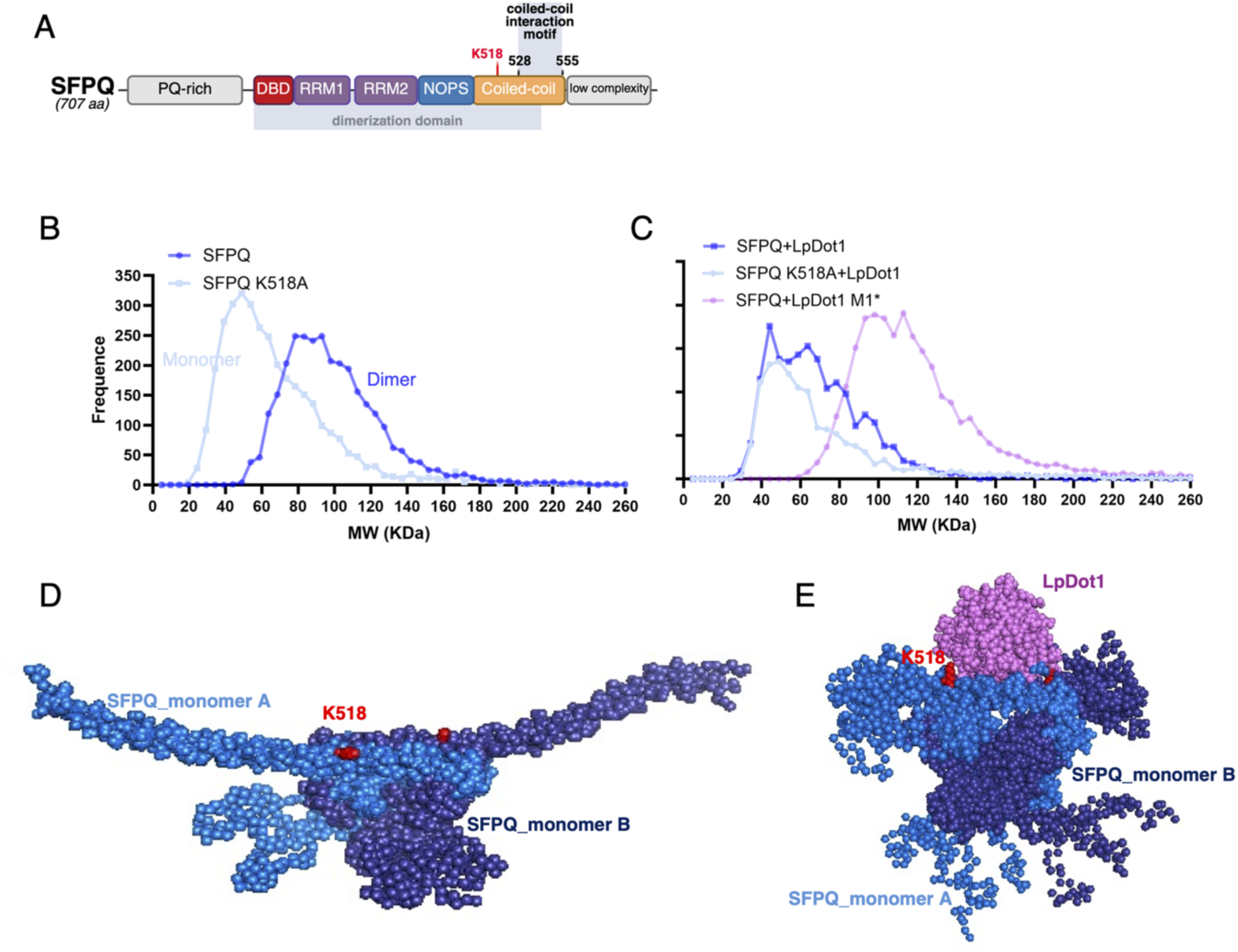
LpDot1 promotes structural rearrangement of the SFPQ coil-coiled impairing its dimerization. (**A**) Schematic representation of human SFPQ domains (according to ^31^). PQ: Pro/Gln rich domain; DBD: DNA-binding domain; RRMs: RNA recognition motifs; NOPS: NONO/ParaSpeckle domain. (**B-C**) Mass photometry analyses. The x-axis displays the measured molecular weight (kDa) and the y-axis the frequency of each observed mass. All data is the results of three independent biological replicates. SFPQ and LpDot1 monomer theoretical molecular weight are 41.5 and 25.7 kDa, respectively. (**B**) Mass distribution of SFPQ vs SFPQ-K518A showing a dimerization peak at 85 kDa and a monomeric peak at 45 kDa, respectively. (**C**) Analysis of complexes formed by SFPQ or SFPQ-K518A with the LpDot1 or the catalytic mutant LpDot1-M1* and the SAM methyl donor. The incubation of SFPQ and SFPQ-K518A with LpDot1 give a profile corresponding to the monomeric peak of SFPQ, whereas the combination of SFPQ with LpDot1-M1* yields a major peak near 100-120 kDa, suggesting a dimeric configuration of SFPQ. (**D**) Best model of SFPQ in solution generated using CORAL^34^. Monomer A and B of SFPQ are represented in light and dark blue, respectively. K518 is in red. (**E**) Best model of LpDot1-SFPQ complex in solution generated by CORAL. SFPQ is represented as in (E) and LpDot1 in purple.

To investigate the role of K518 in SFPQ dimerization, we used mass photometry and assessed the degree of oligomerization of wild-type and mutant SFPQ K518A. In the presence of SAM, SFPQ exhibits an oligomeric mixture dominated by dimers appearing as a single peak (85 kDa). Introduction of the alanine substitution at the 518-position strongly disturbed the monomer dimer equilibrium and reduced dimer formation, resulting in the accumulation of the monomeric form (45 kDa) (**Fig. 3B**). To rule out the possibility that the monomeric state of SFPQ K518A was due to overall protein destabilization, we performed circular dichroism analysis. The resulting spectra were similar for both proteins, indicating that the monomerization was not the result of protein unfolding (**Fig. S4B**). Thus, the K518 residue plays an essential role in SFPQ oligomerization.

We then analyzed the impact of K518 methylation on the oligomeric state of SFPQ by mass photometry analyses of SFPQ or SFPQ K518A incubated with LpDot1 and SAM. We observed that both SFPQ + LpDot1 and SFPQ K518A + LpDot1 conditions display a mass shift around 60-80 kDa, compatible with the complex of monomeric SFPQ and LpDot1 (**Fig. 3C**). Interestingly, when we incubated SFPQ and the catalytically inactive LpDot1-M1*, we recorded a mass peak at 100-120 kDa corresponding to the dimeric form of SFPQ bound to LpDot1 (**Fig. 3C**). These findings identify a critical role of K518 in maintaining SFPQ oligomerization as both K518 substitution and methylation disrupt this process and lead to a shift towards a monomeric state.

### K518 methylation triggers the disassembly of the SFPQ coiled-coiled arms

To provide structural evidence of SFPQ monomerization induced by methylation of K518, we examined the stoichiometry and kinetics of the formation of the SFPQ-LpDot1 complex. We verified that recombinant SFPQ adopts the expected dimeric architecture in solution (**Fig. S4C-G**). The SEC-SAXS profile indeed corresponds to a dimer (R_g_ = 51 Å, D_max_ = 189 Å; **Fig. S4C**) with a folded core and extended flexible segments, as confirmed by the dimensionless Kratky plot (**Fig. S4D-E**). We used the published extended structure of SFPQ (PDB 4WIJ^31^) together with the crystal core (PDB 7UK1^33^) and rebuilt the missing flexible regions with CORAL^34^, obtaining a best-fit model (χ² = 2.22; **Fig. S4F–G**) that captures the elongated coiled-coil and its interaction interface.

The determination of the SFPQ – LpDot1 complex formation kinetics revealed rapid association/dissociation k_on_ / k_off_ constants (**Fig. S5A**) and a weak equilibrium affinity (Kd ≈ 30–50 µM), consistent with an extremely transient complex. Mass photometry showed that even at low concentrations (4 nM), the complex forms quickly (within minutes) but remains only as a minor population (28 % of [1:1] and 12% of [2:1] SFPQ - LpDot1 complexes detected) (**Fig. S5B**). To elucidate further the structure of the complex, we performed SAXS at elevated concentrations of LpDot1 (225 µM). The resulting pair-distribution function yielded R_g_ = 38 Å and D_max_ = 130 Å, compatible with a compact [2:1] SFPQ - LpDot1 complex (**Fig. S5C-D**). The Kratky plot confirmed that the complex is more compact than SFPQ alone (**Fig. S5E**) and a reliable model could be built using the core crystal structure (PDB 7UK1) and rebuilding the flexible segments with CORAL^34^, which nicely fitted the SAXS data (χ² = 2.22; **Fig. S5F**). According to this model, the binding of LpDot1 disrupted the long antiparallel coiled-coil region of SFPQ (present in SFPQ alone, **Fig. 3D**), partially unwinding the corresponding α-helices and bringing the core into a more compact arrangement (**Fig. 3E** and **S5G**). This structural rearrangement correlates well with the shift towards a monomeric population observed upon methylation of SFPQ *via* LpDot1. Thus, our results provide a mechanistic link between the enzymatic activity of LpDot1 and the destabilisation of the SFPQ dimers.

### LpDot1 acts as a *L. pneumophila* virulence factor that affects paraspeckle composition during infection

To analyze whether LpDot1 is important for intracellular replication of *L. pneumophila*, we generated a *lpdot1* deletion mutant (*Δlpdot1*) and compared its survival in host cells to that of the wild-type (WT) *L. pneumophila* strain. Bacterial growth in liquid media was not affected (**Fig. S6A**), but replication of the *Δlpdot1* mutant in the human monocytic cell line THP-1 was significantly delayed as compared to the WT strain (**Fig. 4A**). Complementation of the *Δlpdot1* mutant with a plasmid carrying the *lpdot1* gene under the control of its own promoter restored the ability of *L. pneumophila* to infect eukaryotic cells to a similar level as the WT strain (**Fig. 4B**). Similarly, in human primary macrophages (hMDMs) the *Δlpdot1* mutant displayed a marked replication defect that was fully restored by complementation (**Fig. 4C** and **Fig. 4D**). LpDot1 contributed also to intracellular survival at late stages of infection in its environmental host *A. castellanii* (**Fig. S6B**). Thus, LpDot1 acts as a virulence factor and contributes to *L. pneumophila* survival within human host cells.

**Figure 4.**
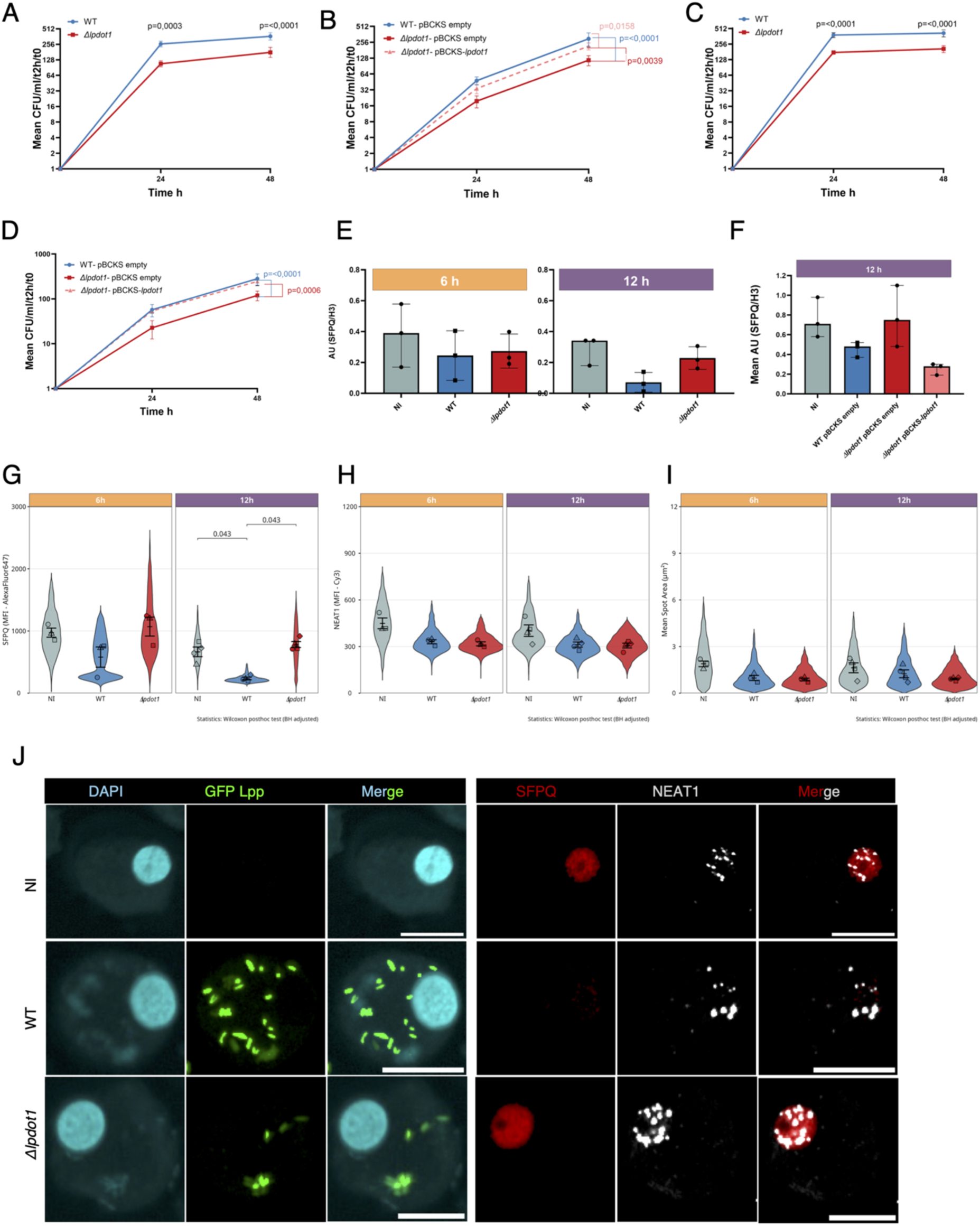
LpDot1 is a virulence factor that affects paraspeckle composition in the host. (**A** and **B**): Intracellular replication of *L. pneumophila* WT and *Δlpdot1* mutant (**A**) or complemented strains (**B**) in THP-1 at a multiplicity of infection (MOI) of 10 was determined by recording the number of colony forming units (CFU) through plating on buffered charcoal yeast extract (BCYE) agar (log2 ratio CFU/ml/t2h/t0). Results are reported as a mean ± SD (n = 3). (**C** and **D**) as (**A** and **B**), respectively but for hMDMs. Differences between groups were analysed using two-way ANOVA with Tukey’s post hoc test (p-values). CFU/ml/t2h/t0 values are presented as mean ± SD. (**E** and **F**) Analysis of the endogenous levels of SFPQ in THP-1 cells by Western-blot, normalized for histone H3 expression (SFPQ/H3) and expressed as Arbitrary Units AU. Results are reported as a mean ± SD of AU, n=3. (**E**) Uninfected (NI) THP-1 cells compared to cells infected with either *L. pneumophila* WT or *Δlpdot1* mutant for 6- and 12-hours. (**F**) Cells were infected for 12 h with WT strain carrying empty pBS-KS or *Δlpdot1* mutant carrying either the empty plasmid or pBCKS-*lpdot1.* (**G-I**) Single cell analysis of paraspeckles in hMDMs left uninfected (NI) or infected with either GFP-expressing *L. pneumophila* WT and *Δlpdot1* (MOI=10), for 6- and 12 hours. (**G**) Violin plots displaying the distribution of AlexaFluo647 fluorescence intensity (SFPQ levels) measurements within the nucleus. (**H**) Violin plots displaying the distribution of Cy3 fluorescence intensity (NEAT1 levels) measurements within the nucleus. (**I**) Violin plots displaying the distribution of NEAT1 area (paraspeckle size) measurements within the nucleus. Each donor is represented by a symbol, and we performed donor-level statistics, calculated as mean ± SD across all single cells from the same donor. Statistical comparisons across donors were performed using the Wilcoxon–Mann–Whitney post hoc tests (BH-adjusted p-value). (**J**) Representative images corresponding to hMDM cells infected 12 h. Nucleus: cyan (DAPI), *Legionella:* green, (GFP), SFPQ: red (Alexa647), and NEAT1: white (Cy3). Scale bar 20 μm.

To understand the mechanism by which LpDot1 contributes to *L. pneumophila* infection, we hypothesized that its direct effect on SFPQ dimerization may alter SFPQ functions, thereby impacting paraspeckle content as well as the downstream cellular processes they regulate. Indeed, SFPQ dimerization is essential for paraspeckle formation and functions, such as the rapid and reversible association and dissociation to its protein partners as well as the correct binding and stabilization of NEAT1, the long non-coding RNA that builds the structural scaffold of paraspeckles^35^. When we monitored SFPQ levels in THP-1 cells during *L. pneumophila* infection, we recorded a decrease of its endogenous levels at late time point of infection (12 h) in a LpDot1-dependent manner (**Fig. 4E**). Complementation of the mutant with the endogenous *lpdot1* gene restored SFPQ levels similarly to those observed for the WT strain. (**Fig. 4F**). Thus, *L. pneumophila* infection has a previously unrecognized global effect on paraspeckle composition that is dependent on LpDot1 secretion.

To further investigate the effect of *L. pneumophila* infection on paraspeckles, we used RNA-FISH coupled to immunofluorescence to monitor in parallel the presence of the SFPQ protein and of the lncRNA NEAT1 in the same cell during infection with the *L. pneumophila* WT or the *Δlpdot1* mutant strains. We quantified the intensity levels of nuclear SFPQ and NEAT1 and assessed paraspeckle morphology by measuring their area (**Fig. S6C**). Indeed, under various stress conditions, paraspeckles can transition to larger, dynamic condensates, modulating the sequestration of specific proteins and RNA, thereby regulating gene expression to overcome the stress^35^. Using this approach, we confirmed the LpDot1-dependend decrease of SFPQ levels within THP-1 cells at late time points of *L. pneumophila* infection (**Fig S6D**). However, no changes neither in NEAT1 levels nor in the size of paraspeckles were observed (**Fig. S6E** and **Fig. S6F**). Single-cell RNA-FISH analysis in hMDMs revealed a faster LpDot1-dependent reduction of the SFPQ protein content compared to THP-1 cells, starting already at 6 h of infection, followed by a pronounced decrease at 12 h (**Fig. 4G** and **J**), an effect likely linked to the higher infection rate recorded in hMDMs compared to THP-1 cells (**Fig. S6G**). Additionally, we observed reduced levels (∼1.3-fold) of NEAT1 (**Fig. 4H** and **4J)** and paraspeckle spot size (**Fig. 4I**) in both WT and *Δlpdot1* infection conditions, suggesting an infection-dependent rather than *lpdot1*-specific effect. Collectively, these results indicate that *L. pneumophila* infection alters paraspeckle composition primarily through depletion of SFPQ protein levels, without disrupting paraspeckle assembly.

### *L. pneumophila* infection modulates alternative splicing

SFPQ is a multifunctional factor essential for the accurate splicing of long introns. Its loss leads to widespread intron retention^36^ and to repression of cryptic last exons, thereby causing premature transcription termination^37^. Depending on its interacting partners, it can either promote exon inclusion or exon skipping^38^ ^39^. As LpDot1has a major effect on the cellular content of SFPQ, we wondered whether it impacts alternative splicing in the host cell. We performed RNA-seq experiments on THP-1 cells at early and late time points of infection with either the WT or the *Δlpdot1* mutant strain. GFP-expressing bacteria were used to allow FACS sorting of infected cells to exclude noise background of transcripts from uninfected cells. We analyzed short read RNA-seq data using the MAJIQ software package^40^ and determined statistically significant local splicing variations (LSVs) across conditions. A schematic representation of the experimental set-up is shown in **Fig. S7A**. Occurrence of relative changes in alternative splicing from one condition to another were quantified as the difference in percent spliced-in (PSI), named delta PSI (dPSI) (dPSI > 10% and 95% probability of change).

When compared wt infected to non-infected (NI) cells, we observed a consistent pattern of alternative splicing changes during *L. pneumophila* infection already at early time points of infection. We recorded a relatively constant number of LSVs at 4 and 12 h (1306 and 1738), corresponding to 897 and 1122 genes, respectively (**Fig. S7B**). The most frequent events are intron retentions (IRs) and alternative first exon (**Fig 5A**). Moreover, we recorded 36% and 32% *de novo* alternative splicing events at 4 h and 12 h, respectively, revealing that *L. pneumophila* manipulates the host splicing machinery in a unique way. GO analysis of the alternatively spliced genes at the later time point of infection highlighted functions related to RNA splicing itself (**Fig. 5B**) and biological processes such as RNA localization, telomere regulation, chromosome organization and microRNA processing^41,42,43,10^, pointing to a broad reprogramming of nuclear processes. **Fig. 5C** shows representative sashimi plots of the alternative splicing events observed for the *POLR2J2* (tentatively linked to the Pol II machinery) and the *SRSF3* (multifunctional RNA-binding protein that controls splicing, mRNA export, stability, and translation) genes at 4 and 12 h, respectively.

**Figure 5.**
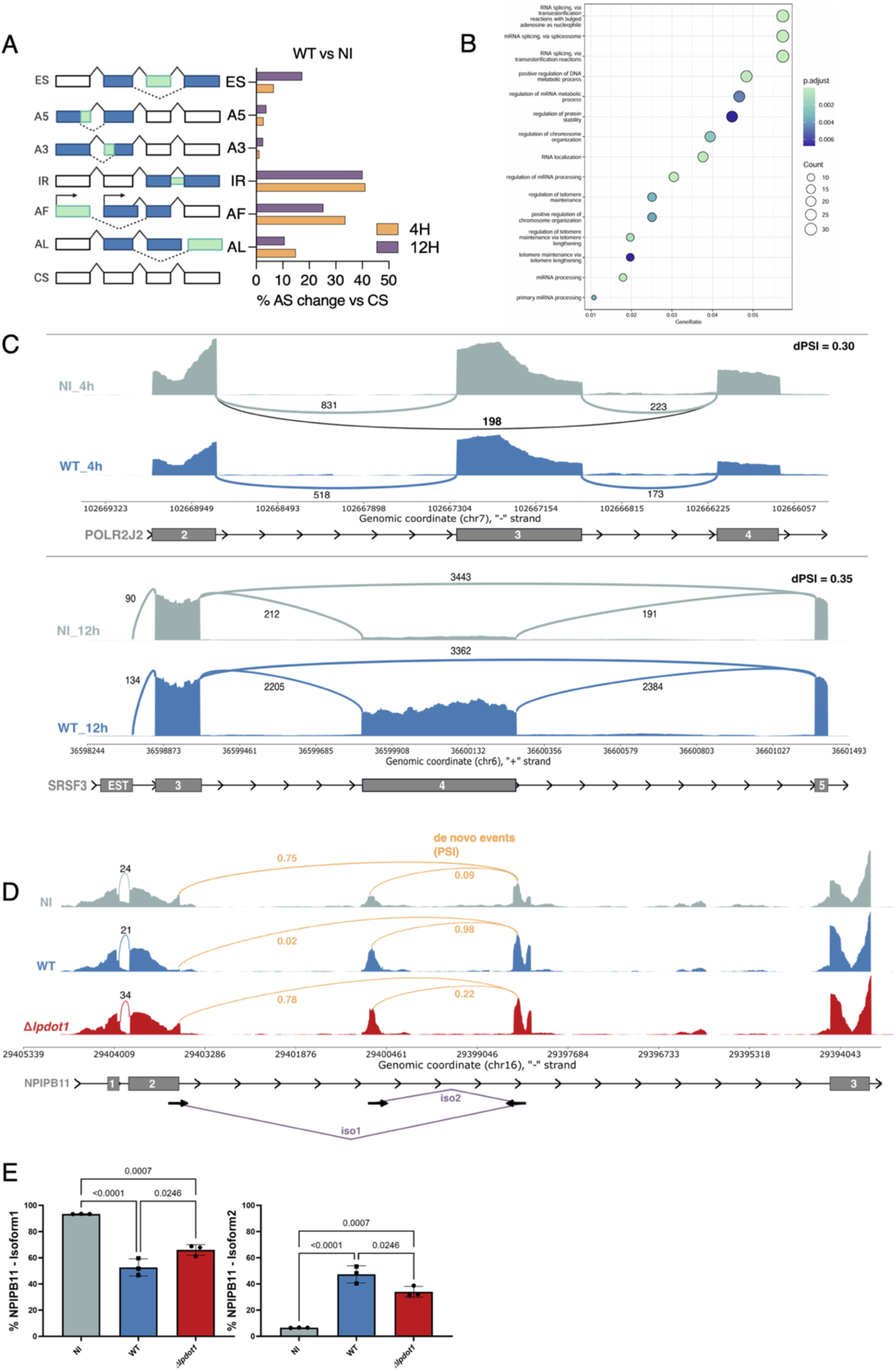
*L. pneumophila* infection modulates alternative splicing. (**A**) *Left*: Schematic representation of the six types of alternative splicing events analyzed: exon skipping (ES), alternative 5’ splice site (A5), alternative 3’ splice site (A3), intron retention (IR), alternative first exon (AF), and alternative last exon (AL). (CS: constitutive splining). Solid lines, constitutive splicing; dashed line, alternative splicing. *Right*: Percentage of change of significant alternative splicing events per each type of event between NI and WT-infected cells at 4 and 12 h. (**B**) Enrichment Gene Ontology (GO) plot showing the top 15 most significantly enriched biological processes for genes showing differences in alternative splicing between NI and WT-infected cells at 12 h (probability of changing > 0.95; all dPSI beyond this threshold were >0.1). The color represents the p-values relative to the other displayed terms (darker blue is more significant) and the size of the terms represents the number of genes that are significant from our list. (**C**) Sashimi plots of two differentially alternative spliced genes, as example. Alternative spliced regions of NI (grey), *L. pneumophila* WT (blue). Per-base expression (transcription intensity) is plotted on y-axis, genomic coordinated on x-axis, and mRNA isoform shown on the bottom (exons in grey, introns as lines with arrow heads). In each plot, a sashimi-like region indicates a heavily transcribed region, in this case exon region. The blank regions between exon regions indicate intronic regions. The arcs crossing exons indicate junction reads. The numbers of junction reads are shown. On the right it displays the estimated dPSI absolute value. (**D**) Sashimi plot of *NPIPB11*. Schematic representation of the *de novo* splicing events occurring in *NPIPB11* gene in orange. Psi values for each junction and condition are indicated. Paired arrows indicate primer sets to validate the *de novo* identified events for Isoform-1 (iso1) and Isoform-2 (iso2). (**E**) qRT-PCR results showing the relative expression of Isoform-1 and Isoform-2, normalized to the housekeeping gene *RPLP0*. Data represent the mean of 3 biological replicates, with each donor indicated by a distinct symbol. Normality was assessed using the Shapiro–Wilk test, and statistical comparisons were performed using ordinary one-way ANOVA.

To assess the role of LpDot1, we compared alternative splicing events of WT vs *Δlpdot1* infected cells. We recorded 49 LSVs (spanning 38 genes) at 4 h and 72 LSVs (spanning 60 genes) at 12 h (**Fig. S7C**), indicating that although most differentially spliced genes are regulated in an infection-dependent manner, a subset of these alternative splicing events strictly depends on the presence of LpDot1. The most frequent event observed is intron retention regardless of the time point of infection (**Fig. S7D**). Furthermore, we identified a large proportion of *de novo* splicing events with 55% and 44% at 4 h and 12 h, respectively. To illustrate *de novo* alternative splicing events, we experimentally validated *NPIPB11* (Nuclear Pore Complex Interacting Protein Family Member B11), a poorly characterized gene with a potential role in nuclear transport. The *NPIPB11* locus showed the highest ΔPSI containing two *de novo* events detected between exons 2 and 3: an alternative 5′ splice site (A5; Isoform-1) and an alternative first exon (AF; Isoform-2) (**Fig. 5D**). These splicing changes were validated by RT-qPCR confirming that isoform-1 levels decreased, whereas isoform-2 levels increased, in an LpDot1-dependent manner (**Fig. 5E**).

### LpDot1 controls the splicing of key genes of the innate immune response favoring immune evasion

Given the profound impact of *L. pneumophila* infection on the host splicing machinery, we investigated whether LpDot1-mediated alternative splicing events could directly impact infection-related genes. Strikingly, we observed *de novo* alternative splicing of *NF-κB2*, one of the five members of the NF-κB family, known to regulate several crucial biological processes such as cell survival and immune responses ^44^. Our RNA-seq data revealed different alternative splicing events on *NF-κB2* between exon 14 and exon 15 (**Fig. 6A**). Besides a canonical exon 14 - exon 15 splicing (Isoform-1), we observed a *de novo* partial intron retention upstream exon 15 (Isoform-2). We measured the differential expression of the two isoforms also by RT-qPCR comparing NI to WT or *Δlpdot1* infected cells and confirmed our observation that WT *L. pneumophila* promotes Isoform-2 generation (**Fig. 6B**). As this is a *de novo* event, we do not know the function of the increase of Isoform-2 confers. Interestingly, exon-15 resides in a region that, when translated, corresponds to the C-terminal portion crucial for its processing and efficient translocation in the nucleus, an important step for triggering inflammation. Thus, to assess possible functional consequences of the increased production of Isoform-2, we quantified the nuclear translocation of NF-κB2 in *L. pneumophila* infected primary monocytes. By using immunofluorescence assays we compared the cytoplasmic and nuclear levels of NF-κB2 during infection and observed significantly less nuclear translocation during WT infection as compared to infection with the *Δlpdot1* strain (**Fig. 6C**). Hence, the higher percentage of the Isoform-2 in WT-infected cells correlates with a dampened immune activation during WT *L. pneumophila* infection, compared to uninfected and *Δlpdot1* infected cells.

**Figure 6.**
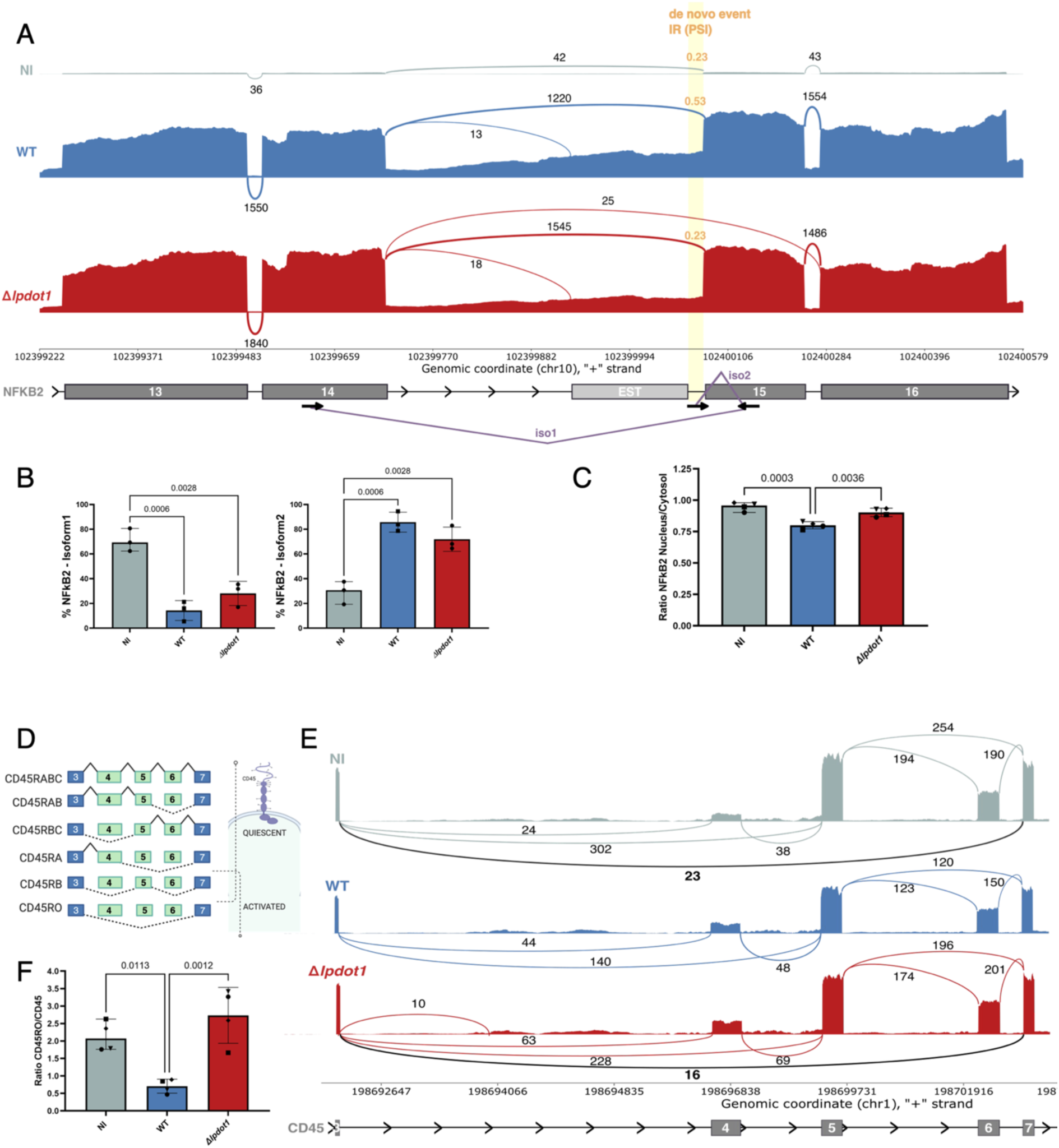
LpDot1 controls alternative splicing of key genes of the innate immune response favoring immune evasion. (**A**) Sashimi plot of *NFKB2*. Schematic representation of the *de novo* splicing events occurring in *NFkB2* gene in orange. Psi values for *de novo* IR junction of each condition are indicated. Paired arrows indicate primer sets for the two isoforms (iso1 and iso2) (**B**) qRT-PCR results showing the relative expression of Isoform-1 and Isoform-2 for *NFkB2*, normalized to the housekeeping gene *RPLP0*. Data represent the mean of 3 biological replicates, with each donor indicated by a distinct symbol. Normality was assessed using the Shapiro–Wilk test, and statistical comparisons were performed using ordinary one-way ANOVA. (**C)** Immunofluorescence analysis of NFκB2 nuclear translocation in primary monocytes left uninfected (NI) or infected with either *L. pneumophila* WT or Δlpdot1 GFP-expressing strains (MOI = 10) for 12.5 hours. Nuclear fluorescence intensity of AlexaFluor647 (NFκB2) was divided by its cytoplasmic intensity of the same cell. Data represents the mean of four biological replicates, with each donor indicated by a distinct symbol. Normality was assessed using the Shapiro–Wilk test, and statistical comparisons were performed using ordinary one-way ANOVA. (**D**) Schematic of the variably spliced region of the *CD45* gene and resulting isoforms. Predominantly expressed isoforms in resting (naive) and activated cells are as indicated. (**E**) Sashimi plot of *CD45*. Alternative spliced regions of NI THP-1 (grey) or infected 4 hours with *L. pneumophila* WT (blue) and Δ*lpdot1* mutant (red) strains. Per-base expression (transcription intensity) is plotted on y-axis, genomic coordinated on x-axis, and mRNA isoform shown on the bottom (exons in grey, introns as lines with arrow heads). (**F**) Analysis of CD45RO levels in primary monocytes left NI or infected with either *L. pneumophila* WT or *Δlpdot1* GFP-expressing strains (MOI = 10) for 5 hours by flow cytometry. CD45RO levels (APC Fire 750 fluorescence intensity) were normalized to total CD45 levels (PE fluorescence intensity) (gating strategy in **Fig. S8**). Data represent the mean of four biological replicates, with each donor indicated by a distinct symbol. Normality was assessed using the Shapiro–Wilk test, and statistical comparisons were performed using ordinary one-way ANOVA.

Another interesting alternative splicing event was observed in the *CD45* gene. CD45, or PTPRC (Protein Tyrosine Phosphatase Receptor type-C), is an important regulator in the immune system whose expression is known to be tightly regulated^45^. *CD45* encodes for a transmembrane domain tyrosine phosphatase for which five isoforms that differ in their extracellular domains are generated through alternative splicing of exons 4, 5, and 6 (**Fig. 6D**). Quiescent immune cells predominantly express the largest CD45 isoforms, whereas shorter isoforms are privileged in more activated states^46^. SFPQ has been shown to facilitate the exclusion of all these three exons, a process that leads to the generation of the CD45RO isoform which is expressed on activated immune cells ^47^. Specifically in monocytes, it has been reported that the surface expression of CD45RA and CD45RO is tightly regulated during inflammation, with an increase of the CD45RO isoform upon LPS stimulation^45^. We recorded reduced splice junction reads connecting exon 3 to exon 7 (corresponding to CD45RO isoforms) at 4 h when cells were infected with WT *L. pneumophila* strain, compared to NI or *Δlpdot1* infected cells (**Fig. 6E**). To investigate whether LpDot1 has an impact on the surface expression of the activation state isotype of CD45, we performed flow cytometry and measured the ratio of CD45RO/total CD45 (gating strategy in **Fig. S8**). Consistent with our RNA-seq data, LpDot1 expression led to decreased levels of CD45RO during *L. pneumophila* infection (**Fig. 6F**), likely masking the activated/infected state of the bacteria containing host macrophages from the systemic immune system.

Taken together, our data demonstrate that *L. pneumophila* actively targets the paraspeckle protein SFPQ to manipulate the host splicing machinery to dampen innate immune responses, thereby preserving its replication niche. These findings not only reveal an additional strategy a bacterium may employ to hijack host responses but also shed light on previously unknown regulatory mechanisms of SFPQ and paraspeckles.

## DISCUSSION

In this study, we revealed a previously unrecognized mechanism by which *L. pneumophila* manipulates host nuclear cellular processes to promote infection. Our results show that LpDot1, a bacterial DOT1-like histone methyltransferase, retains the core catalytic architecture of DOT1 MTases, but exhibits a different substrate specificity as it directly methylates the RNA-binding protein SFPQ, a key scaffold within paraspeckle structures. This feature distinguishes LpDot1 from previously characterized bacterial DOT1 homologs. For instance, *Mycobacterium tuberculosis* encodes a DOT1-like methyltransferase (Rv2067c) that keeps the activity of its eukaryotic homologs by trimethylating histone H3K79 in macrophages, and it decreases the DOT1L-mediated epigenetic mark on pro-inflammatory genes^48^. A possible explanation for the functional divergence is that *L. pneumophila* acquired a nucleosome-targeting enzyme, but evolutionary modifications have redirected its substrate specificity from nucleosomal histones to alternative targets aiding bacterial replication. On the other hand, since the targets of the DOT1 homologs in protists are unknown, it is also plausible that *Legionella* may have acquired an enzyme that originally targeted non-histone proteins that later evolved in higher eukaryotes to target nucleosomes, or it still retains this activity in higher eukaryotes too, but this specific target remained yet unidentified.

*L. pneumophila* proteins are examples how a human intracellular pathogen can repurpose such horizontally acquired enzymes to manipulate host cell functions to its advantage^18^. A key example is RomA, which catalyzes the trimethylation of histone H3 at lysine 14 (H3K14) during infection^20^. This epigenetic mark was initially considered novel in the mammalian host, but based on our findings it was recently confirmed to occur endogenously also in human cells^49^, highlighting *Legionella*’s potential as a tool to uncover previously undefined eukaryotic pathways. *dot1* genes seem essential to *Legionella* lifestyle, as they recently arose likely through multiple horizontal gene transfers among prokaryotes, eukaryotes, and viruses sharing the same ecological niche ^25^. This observation supports the hypothesis of a selective pressure favoring the acquisition and retention of the *dot1* genes across *Legionella* spp.

Here we revealed that LpDot1 is an active MTase mainly targeting RNA-binding proteins and paraspeckle components such as SFPQ, but also PCBP1 that was found exclusively methylated by LpDot1, and FUS, HNRNPA1, and HNRNPA2B1 that showed an altered methylation pattern, suggesting a broad effect of *Legionella* infection on paraspeckle composition. It will be exciting to explore these modifications further in the future. Here we further analyzed SFPQ methylations and revealed its function. We detected a previously unrecognized methylation site on K518 of SFPQ, which is located within its coil-coiled region. While several arginine residues of SFPQ are known to be post-translationally modified (*e.g.* by methylation or citrullination), only K314 has previously been reported to be methylated and K518 had been described as SUMOylated (PTMeXchange database)^50^, but we now showed it can also be methylated. We further demonstrated that any perturbation of K518, whether through amino acid substitution or methylation, significantly alters the oligomerization state of SFPQ, which is crucial for its activity^31^ ^32^. Importantly, by studying the LpDot1 – SFPQ complex in solution we were able to propose a mechanistic model by which LpDot1-catalyzed methylation of K518 disrupts the long antiparallel coiled-coil of SFPQ, a crucial region for its oligomerization and quaternary structure-dependent function. We therefore wondered whether this destabilisation could undermine paraspeckle formation, ultimately remodelling the host’s transcriptional landscape in favour of the pathogen.

Paraspeckles are dynamic nuclear bodies involved in regulating gene expression through the sequestration of nascent transcripts and modulating the splicing process. Dynamic assembly and disassembly of paraspeckles has been reported in response to LPS treatment of human macrophages to likely orchestrate the immune response against bacterial pathogens^14^. We showed that SFPQ methylation by LpDot1 results in a substantial decrease of its cellular levels during *L. pneumophila* infection. However, despite the drastic reduction of SFPQ, we did not observe paraspeckle’s disassembly, as NEAT1 levels and spot sizes showed only a modest reduction. This observation is consistent with previous reports showing that only the NEAT1 middle domain is necessary and sufficient for paraspeckle assembly with NONO and FUS playing an important role in this process^51^ ^52^. Notably, similar perturbations of key paraspeckle components without complete disassembly have been described in viral infections: SFPQ is cleaved by direct proteolysis during rhinovirus infection of human epithelial cells^53^, and FUS levels are markedly reduced in B cells infected with Kaposi’s sarcoma–associated herpesvirus^54^. Together with our data, these findings suggest that pathogens employ a conserved strategy to remodel paraspeckle composition by selectively targeting their core proteins, thereby altering host gene regulation while preserving paraspeckle integrity. We can speculate that the decrease of SFPQ content observed during *L. pneumophila* infection may induce an additional general remodeling of the nuclear architecture, impacting, for example, on speckle dynamics. Indeed, speckles and paraspeckles are spatially adjacent and their interplay and redistribution have been reported in viral infections ^55^. Moreover, it has been reported that SFPQ depletion causes speckles/paraspeckles fusion, suggesting that the paraspeckle shell composition dictates the independence of MLOs in the nucleus, therefore defining their specific features and functions^56^. The impact of bacterial infections on speckle dynamics has been recently explored. It was shown that *Listeria monocytogenes* infection changes the speckle morphology by redistributing the splicing factor SC35, an essential component of nuclear speckles^57^.

The manipulation of the host splicing machinery is emerging as an effective strategy that viral and bacterial pathogens use to modulate the host immune response^57^ ^58^ ^59^, however the targeting of the paraspeckle protein SFPQ to modulate alternative slicing was not reported before. In the current study, we revealed a significant impact on the splicing machinery at early time points of infection by *L. pneumophila*, distinct from other intracellular bacteria such as *Listeria monocytogenes*, which modulates alternative splicing at late time points of infection in a pore-forming toxin dependent manner ^58^. The fast and broad infection-related effect we report here is in line with other works showing an LPS-induced impact on alternative splicing^60^ ^61^ ^62^. Interestingly, more than one third of splicing events were *de novo* events, suggesting that *L. pneumophila* infection subverts the host splicing machinery in a unique way. GO analyses revealed that the genes that are alternatively spliced during infection are enriched in biological processes directly related to alternative splicing regulation *per se,* RNA localization, telomere regulation, chromosome organization and microRNA processing, indicating a broad reprogramming of nuclear processes. A possible role of LpDot1 activity in directly influencing the splicing machinery is supported by STRING analyses of the LpDot1 - mediated methylation targets identified by MS/MS as they comprise a cluster of 21 proteins that is enriched for spliceosomal components (**Fig. S9A**). Furthermore, our RNA-seq data revealed a specific subset of LpDot1-dependent differentially spliced genes that are linked to lipid metabolism, osmotic stress response and mitophagy (**Fig. S9B**). This suggests that modulation of alternative splicing may also play a regulatory role in membrane remodeling and key stress-responses and a control of mitochondrial functions, limit damage, or modulate immune responses.

Importantly, we functionally validated alternative spliced events in infection-related genes. LpDot1 favors a specific intron retention upstream exon-15 of *NF-κB2.* NF-κB2 is a player of the noncanonical NF-κB pathway, which many viruses and bacteria including *L. pneumophila* have been shown to manipulate, mainly by post-translational modifications of proteins or inhibition of translation ^63^ ^64^ ^65–68^. NF-κB2 is produced as an inactive precursor, p100, that undergoes proteolytic processing to yield the functional subunit p52 that subsequently migrates to the nucleus to regulate target genes. Recent studies reported alternative splicing events of *NF-κB* genes, generating several transcripts corresponding to alternative isoforms. In mice for example, a splice variant of *NF-κB1* involving skipping of exon 13 results in a 79-nt deletion^69^. In the human Hodgkin lymphoma cell line L-1236, researchers reported temporal changes in the expression ratio of *NF-κB2* isoforms, a phenomenon proposed to contribute to malignant transformation through uncontrolled activation of the non-canonical NF-κB pathway^70^. Mechanistically, such splicing events can alter the balance between the precursor form p100, which retains NF-κB complexes in the cytoplasm, and the processed form p52 that translocates to the nucleus and activates transcription. To our knowledge, this is the first example of a bacterium inducing a variant of NF-κB2 that may be processed less efficiently, leading to reduced p52 and increased p100-like inhibition. Beside *NF-κB2*, we also identified a differentially spliced isoform of NF-KBID, a negative regulator of the canonical NF-kB1 signal transduction, therefore suggesting that LpDot1 is a unique factor that mediates fine tuning of the entire NF-κB axis in an unconventional way.

Along with NF-κB2 regulation, we demonstrated that *L. pneumophila* infection modulates CD45 isoforms expression, leading to a shift in the processing of CD45 pre-mRNA that results in expression of the smaller CD45 isoforms (CD45RO), a commonly recognized hallmark of cell activation. The function of CD45 is extensively characterized in T-cells, as it is essential for antigen receptor signaling and proper immune function, but it has also been shown to be important in monocyte and macrophage innate immune responses^45–47^. A direct link between SFPQ and CD45 splicing has been reported in T-cell activation when the recruitment of SFPQ to the CD45 pre-mRNA facilitates CD45RO isoform at the cell surface^47^. However, current evidence does not document a direct involvement of SFPQ in CD45 splicing in monocytes/macrophages.

Here we shed light on the essential role of paraspeckles and the host splicing machinery system in controlling infections and show how *L. pneumophila* can be used as a tool to uncover and understand yet uncharacterized eukaryotic pathways. Our findings may also pave the way for new antibacterial strategies. The 2.1 Å resolution of LpDot1 structure provides exceptional detail for structure-based drug design. The clear SAH binding mode reveals multiple contact points for inhibitor development, while the structural differences between LpDot1 and human DOT1L offer opportunities for selective targeting. Key druggable features include distinct substrate-binding loops that could be targeted by allosteric inhibitors; altered electrostatic surfaces enabling selectivity over human enzymes; and unique conformational flexibility in substrate-recognition regions. Such selective inhibitors could represent a novel class of anti-virulence therapeutics that disrupt *Legionella*’s ability to manipulate host nuclear processes without directly targeting bacterial viability, potentially reducing resistance development.

## RESOURCE AVAILABILITY

Further information and requests for resources and reagents should be directed to and will be fulfilled by the lead contact, Monica Rolando.

## MATERIAL AVAILABILITY

This study did not generate new unique reagents.

## DATA AND CODE AVAILABILITY

The raw sequencing data generated in this study are deposited in https://www.ncbi.nlm.nih.gov/geo/ (accession number pending). The mass spectrometry proteomics raw data have been deposited in the ProteomeXchange Consortium via the PRIDE partner repository. Data are available via the identifier PXD071877. The final refined model of the LpDot1 structure was deposited with wwPDB ID code pdb_00009NE1. Any additional information required to reanalyze the data reported in this paper is available from the lead contact upon request.

## COMPETING INTERESTS

The authors declare no competing interests

## ACKNOWLEDGMENTS

We thank Laura Gomez Valero and Pierre Foucault for fruitful discussions and technical help. Work in the C.B. laboratory was supported by the “Fondation de la Recherche Médicale” grant EQU202503020066 to CB. the PTR (Programmes Transversaux de Recherche) grant PTR 395-20 from Institut Pasteur Paris and ANR-25-CE15-6707 to M.R. S.N. was funded by PTR 395-20, “Agence Nationale de Recherche” grants ANR-10-LABX-62-IBEID and ANR 20-PAMR-0011-TheraEPI. H.S. was funded by ANR-18-CE15-0005-01 to M.R. J.E.M. was funded by ANR-10-LABX-62-IBEID. We gratefully acknowledge the support of the Swing Beamline (Synchrotron SOLEIL, Saint Aubin) and Aurélien Thureau for SAXS experiments and the C2RT core facilities at the Institut Pasteur, in particular Elodie Turc, Laure Lemée, and Rania Ouazahrou from the Biomics Facility supported by France Génomique (ANR-10-INBS-09) and IBISA; Stéphane Petres and Mireille Nowakowski of the “Production and purification of recombinant proteins” Technological platform; Patrick England and Sébastien Brulé from the Molecular Biophysics platform, Magalie Duchateau (Proteomics platform) for contributing to the initial MS analysis, Christian Weber and Anne Danckaert from the Photonic BioImaging platform (UTechS PBI) supported by the French National Research Agency (France BioImaging, ANR-24-INBS-0005 FBI (BIOGEN); Investments for the Future), the Institut Pasteur and the Région Île-de-France (DIM1Health program) funding for the use of the Opera Phenix system.

## AUTOR CONTRIBUTIONS

Conceptualization: S.N., C.B. and M.R. Methodology: S.N., S.M., N.M., B.R., R.E., J.D.R., N.L., J.E.M., H.S., S.S. Bioinformatical analyses C.R. and Q.G.G. Analysis and interpretation of the data: S.N., S.M., C.R., B.R., R.E., Q.G.G., M.M., A.B., C.B., and M.R. Writing – original draft: S.N., M.R., C.B. Writing – review and editing: S.N., M.R., C.B., S.M., M.M, and A.B. Supervision and funding acquisition: M.R., C.B., and A.B.

## MATERIALS and METHODS

### Bacterial strains and growth conditions

*Legionella pneumophila* strain Paris was grown on ACES-buffered charcoal yeast-extract (BCYE) agar and on (ACES)-buffered yeast extract broth (BYE) at 37 °C^71^. For the generation of *lpdot1* knock out, approximately 500 bp of sequence upstream and downstream of *lpp3025* was used to flank an Apramycin resistance cassette and *mazF* (aprR-mazF). Fragments were amplified by PCR (primers IDs 15-18, **Table S2**), the aprR-mazF was amplified from pJM028 using (primers IDs 68-69, **Table S2**). Each of the three PCR fragments were then ligated into pGEM T easy (PROMEGA) cut with *Not*I-HF (Thermofisher) using NEBuilder HiFi master mix (New England Biolabs). The resulting plasmid was checked by WPS, transformed into *E. coli* DH5α, then used as the template to generate approximately 1 µg of linear DNA by PCR, which was transformed as previously described^21^. Strains were checked by PCR and sequencing (primers IDs 70-71, **Table S2)** to check the deletion mutant. The *Δlpdot1* mutant was complemented by cloning into pBCKS (Stratagene) the full-length *lpdot1* gene with an N-terminal myc-tag under the control of its own promoter. Wild-type (WT) and *Δlpdot1* mutant strains expressing EGFP were obtained as previously described^21^. Bacterial growth was monitored in broth cultures using a Tecan plate reader. Optical density at 600 nm (OD₆₀₀) was measured every 20 min for 40 h at 37 °C, and growth curves were generated from the recorded absorbance values. For *Escherichia coli* Luria-Bertani broth (LB) was used. *E. coli* strain DH5α cells were used for DNA cloning purposes, transformed according to standard methods. The *E. coli* BL21(DE3) pLysS and Rosetta2 (DE3) (Merck Millipore) strains were used for protein expression. When needed antibiotics were added: for *L. pneumophila* (*E. coli*): kanamycin 12.5 μg/ml (50 μg/ml), gentamycin 12.5 μg/ml, apramycin 15 μg/ml, chloramphenicol 10 μg/ml (10 μg/ml), and ampicillin (only for *E. coli*) 100 μg/ml.

### Cell culture

The human monocyte cell line THP-1 (ATCC TIB-202TM) was maintained in RPMI GlutaMAX supplemented with 10% fetal bovine serum (FBS, Biowest) at 37 °C in a 5% CO₂ humidified atmosphere. When stated, undifferentiated THP-1 cells were seeded in RPMI containing 50 µg/ml phorbol 12-myristate 13-acetate (PMA) to induce differentiation into adherent cells (dTHP-1) for 3 days. Human blood was collected from healthy donors under the ethical rules established by the French National Blood Service (EFS). Peripheral blood mononuclear cells (PBMCs) were isolated by Ficoll-Paque PLUS density-gradient separation (density 1.077 mg/ml, Cytiva) at room temperature. PBMCs were incubated with anti-human CD14 antibodies coupled to magnetic beads (Miltenyi Biotec) and subjected to magnetic separation using MS/LS columns (Miltenyi Biotec). Positive selected CD14 + cells were plated in RPMI 1640 Medium, GlutaMAX™ Supplement, HEPES (Life Technologies), and 20% heat-inactivated FBS. Human monocyte-derived macrophages (hMDMs) cells were differentiated with 50 ng/ml of recombinant human macrophage colony-stimulating factor (rhMCSF, Miltenyi Biotec) for 6-7 days at 37°C with 5% CO₂ in a humidified atmosphere. At day 3, additional rhMCSF (50 ng/ml) was added. *Acanthamoeba castellanii* strain C3 (ATCC 50739) trophozoites were grown at 20°C in PYG medium ass described ^72^. Infection buffer was prepared using PYG 712 medium [2% proteose peptone, 0.1% yeast extract, 0.1 M glucose, 4 mM MgSO₄, 0.05 mM Fe(NH₄)₂(SO₄)₂•6H₂O, 2.5 mM NaH₂PO₄, 0.4 mM CaCl₂, 0.1% sodium citrate dihydrate, 2.5 mM K₂HPO₄].

### Infection conditions and replication assays

Infection assays were performed as previously described^21^. For all conditions, *L. pneumophila* strains were prior grown in BYE to stationary phase (OD_600_ ∼ 4.2). THP-1 and hMDM cells were infected with a multiplicity of infection (MOI)=10, *A. castellanii* with MOI=0.1. For replications assays in human cells *L. pneumophila* intracellular replication was recorded after 2-, 24- and 48 h post-infection by enumeration of colony forming unit (CFU) after plating on BCYE. Replication assays in *A. castellanii* trophozoites were performed at 20°C in PYG 712 medium without peptone, glucose and yeast extracts. Bacterial number was determined every 24 h for 7 days by CFU enumeration. Bacteria counting were normalized to the input number (CFU/ml at t 0h), and to the entry number (CFU/ml at 2h). Statistical comparisons across donors were performed using the 2way ANOVA test followed by Tukey’s multiple comparisons. Data represents the mean of three biological replicates.

### Cloning, expression and purification of human SFPQ and *L. pneumophila* LpDot1 proteins

#### Recombinant SFPQ

His-Tag SFPQ was produced from pCDF11-SFPQ (276-598), (Addgene #135437) and SFPQ point mutation K518A was generated by site-directed mutagenesis using the QuikChange Site-Directed Mutagenesis kit (Stratagene) (primers ID 5-6 listed in **Table S2**). SFPQ and SFPQK518A were expressed in Rosetta2 (DE3) (Merck Millipore) and purified using nickel-affinity chromatography followed by size exclusion chromatography. The cells were grown in TB broth with 50 μg/ml spectinomycin at 37°C and induced for 18 h at 18 °C with 0.2 mM isopropyl β-D-1-thiogalactopyranoside (IPTG) at an OD_600_ of 1.8-2.2. Cells were harvested by centrifugation at 18000× g for 15 minutes at 4 °C. The cell pellet was resuspended in buffer A (50 mM Tris–HCl (pH 7.5), 500 mM NaCl, 20 mM imidazole, 10% glycerol, 1 mM DTT + 0.4 mg/ml EDTA-free protease inhibitors), lysed using a Cell Disruptor Constant System TS (1.3 kBar) and clarified by centrifugation at 42000 × g for 1 h at 4 °C. The soluble fraction was then applied to a nickel-chelating column (Hi-Trap HP, GE healthcare). SFPQ was eluted using a gradient of 20–500 mM imidazole. The fractions of interest were then applied to a 16/60 Superdex 200 column and developed with 20 mM Tris–HCl (pH 7.5), 150 mM NaCl + 10% glycerol, 1 mM DTT.

#### Recombinant LpDot1

The *lpDot1* gene (*lpp3025*) from *L. pneumophila* strain Paris was cloned in pET-28(a) (Novagen) and codon-optimized by Genecust company. The optimized gene sequence was PCR-amplified using oligonucleotides IDs 1-2 listed in **Table S2** and cloned into XhoI/XbaI sites of pTEM14^73^ by restriction free-cloning (PSM16), to generate a LpDot1 recombinant protein with a hexa-histidine-HRV3C-tagged site at the N-terminus. Site-directed mutagenesis was performed using the QuikChange Site-Directed Mutagenesis kit (Stratagene) to insert G96R and G98R mutations and generate PSM16M* plasmid that carries LpDot1 M* (primers IDs 3-4 of **Table S2**). PSM16 and PSM16M* were transformed into *E. coli* BL21(DE3) pLysS cells, and transformants were selected on LB agar plates containing 50 μg/ml kanamycin and 34 μg/ml chloramphenicol. Cultures were grown at 37 °C in 1 L TB broth (supplemented with 0.2 mM MgSO_4_, 0.8% glycerol, 50 μg/ml kanamycin and 34 μg/ml chloramphenicol) to OD_600_ ∼ 2. Expression was induced with 0.2mM IPTG and continued overnight at 18°C. Cells were harvested by centrifugation and resuspended in lysis buffer [50 mM Tris pH8, 500 mM NaCl, 20 mM imidazole, 5U/ml DNAse, 1 mg/ml hen egg white lysozyme and complete EDTA-free protease inhibitor cocktail (Roche #48679800)]. Following French press lysis and centrifugation, the clarified supernatant was subjected to Ni-NTA agarose chromatography (Cytiva). The resin was washed with 13% Buffer B (50 mM Tris pH8, 500 mM NaCl, 500 mM imidazole), and His_6_LpDot1 was eluted with 100% Buffer B. Final purification was achieved by size-exclusion chromatography (Superdex 75, GE Healthcare) pre-equilibrated in SEC buffer (20 mM Tris pH 7.8, 150 mM NaCl) and run isocratically at 0.8 ml/min (Akta Purifier, GE). Pure His_6_LpDot1 was concentrated to 2.75 mg/ml using Vivaspin 20 centrifugal devices. Protein purity was verified by SDS-PAGE and mass spectrometry. Purified protein was stored in aliquots at -80°C.

### LpDot1 structure determination

#### Crystallization and data collection

His_6_LpDot1 (2.75 mg/ml) was crystallized at 20°C, in complex with SAM (Sigma #A4377) at 1:6 molar stoichiometry (protein:SAM). Crystallization drops (2 μl protein:SAM mixture + 1μl reservoir solution) were set up using hanging-drop vapor diffusion with 1 ml reservoir volume. The optimized reservoir solution contained 0.1 M HEPES pH 8.0, 10 mM TCEP, and 15% PEG 3350. For cryoprotection, crystals were transferred to reservoir solution supplemented with 25% (v/v) glycerol and 0.7 mM SAM, mounted in cryo-loops (Hampton Research), and flash-vitrified in liquid nitrogen. X-ray diffraction data were collected at -107°K at the Protein Crystallography Facility (Institut Pasteur de Montevideo, Uruguay), equipped with a MicroMax-007 HF rotating Cu anode (Rigaku) and a Mar345 image plate detector (marXperts).

#### Structure determination and refinement

Bragg diffraction intensities were indexed and integrated with XDS^74^ and scaled and reduced to amplitudes with Aimless and Ctruncate^75^. The structure was solved by molecular replacement using Phaser^76^ with an AlphaFold2-predicted model of LpDot1 as the search probe. Structure refinement was performed using phenix.refine^76^, iterating with interactive model rebuilding in Coot^77^. Validation was performed with Molprobity^78^ and wwPDB validation tools. The final refined model was deposited with wwPDB ID code pdb_00009NE1, comprising protein residues 34-236 (relative to UniProt reference sequence A0A3A6VYM1), one SAH molecule, and 44 ordered water molecules. In the crystal structure, LpDot1 Lys40 was found to be monomethylated on its sidechain amine.

#### SAXS analysis of LpDot1, SFPQ and SFPQ-LpDot1 complex

SAXS data were collected on the SWING beamline at Synchrotron Soleil (France) using the online HPLC system. The column (Superdex 200 increase 5/150 mm) was equilibrated in buffer containing 20 mM Tris pH 7.8, 150 mM NaCl, 32 µM SAM, 5% glycerol. Samples (50µl of 225 µM of LpDot1, 50µl SFPQ at 52 µM, 50µl of a mix of SFPQ at 52 µM LpDot1 at 225 µM) were injected into a size exclusion column (Superdex 200 increase 5/150 mm) cooled at 15 °C eluting directly into the SAXS flow-through capillary cell at a flow rate of 200 µl min^−1^. The data were analysed using FOXTROT and PRIMUS from ATSAS 3.2^79^, from which Guinier was generated. Scattering curves were selected for stable *R*_g_ at the apex of the elution profile, the selected curves were averaged, and buffer signal was subtracted. The scattering curve of the complex was extracted from the main peak using LC and Evolving Factor analysis in RAW 2.3.1 ^80^. From these corrected scattering curves, the pair distribution function was computed using GNOM (version 5.0)^81^, and the normalized Kratky plot was generated. Using the structure of LpDot1and the alphafold structural model of the complex, the experimental curves were compared to the theoretical curves using CRYSOL (version 2.8.3)^81^ Coral models were generated with CORAL (version 1.1)^34^. The SAXS statistics are provided in **Source Data supplementary file**.

### Methyltransferase (MTase) assay

Radioactive methyltransferase (MTase) assay was performed as in ^82^. Briefly, 2 μg of purified LpDot1 or 1 μg of human DOT1L (Active Motif #31474) were incubated with 5 μg of calf thymus histone (CTH, Sigma) or 10 μg of nuclear extracts (NE, EMD Millipore 12-309) and 32 μM SAM ^14^C as a methyl-donor (PerkinElmer NEC3630100C) in buffer containing 50 mM Tris-HCl, pH 8.0, 20 mM KCl, 10 mM MgCl2, 0.02% Triton X-100, 5% glycerol, 1 mM DTT and 0.4 mg/mL EDTA-free protease inhibitor (Mini Protease Inhibitor Cocktail tablets, ThermoFisher). Nuclear extract from *A. castellanii* was obtained as described in ^83^. All reactions were performed at 30 °C for 14 h if not otherwise stated. To validate LpDot1human targets, the predicted catalytic site of LpDot1, and the methylation site of SFPQ, we used 2 μg of LpDot1 and LpDot1 M* enzymes and 1 μg of the following human recombinant proteins as substrates: MGB3 (MedChem, HY-P70199), HSPA1B (MedChem, HY-P73105), SNX3 (abnova, H00008724-P01) SFPQ and SFPQK518A. Reactions were stopped by the addition of SDS loading buffer and incubation of the samples for 5 min at 95 °C. Samples were separated in 8-16% SDS-PAGE and protein size determined using a pre-stained molecular weight marker (ThermoFisher Scientific). The gel was transferred to a PVDF membrane (0.2 µm; Trans-Blot Turbo system, Biorad) and stained with amido black (AB) (Sigma) for the loading control. The membranes were exposed for 24 h and the signal was read using a Typhoon TRIO Variable mode imager, and images were processed using Fiji software (ImageJ v2.9.0/1.53t).

### Mass photometry measurements

The MP experiments were performed at 20°C using the 2MP instrument (Refeyn, Oxford, UK). The samples considered for this experiment: SFPQ, SFPQ K518A, LpDot1 and LpDot1 M*, SFPQ + LpDot1 (molar ratio 1:3), SFPQK518A + LpDot1 (molar ratio 1:3), SFPQ + LpDot1 M* (molar ratio 1:3), were diluted at 40 nM in presence of 32 μM SAM (NEB). 18 µl of the filtered PBS filtered and 2 µl of each sample were loaded into a well of the gasket (final concentration 4 nM). Immediately after the solution was mixed by pipetting, a 1 min video was recorded and processed using the AcquireMP and DiscoverMP softwares, respectively (Refeyn, Oxford, UK). Ratiometric contrast distribution was plotted as Kernel Density Estimates (KDE) or histograms with a 0.0002 bandwidth or binwidth, respectively, and a 0.013 contrast limit. The molecular mass of the protein was calculated from the MP contrast distribution by applying the calibration obtained using bovine serum albumin (BSA).

### Circular dichroism

Far-UV circular dichroism (CD) spectra (190 to 260 nm) were recorded with a Jasco J-1500 spectropolarimeter (Jasco, Tokyo, Japan) with a 450W lamp at 20°C using a 100-µm-pathlength quartz Suprasil cell (31/B/Q/0.1, Starna). SFPQ and SFPQ K518A were resuspended at around 10 and 7 µM respectively, in a buffer containing 50 mM TRIS, 150 mM NaCl, 10% glycerol and 1mM DTT. Spectral acquisitions of 0.1 nm steps with 2 s integration time at 50 nm/min and a bandwidth of 1 nm were performed 4 times for the samples as well as for the buffer. The measurements were carried out with constant nitrogen gas flux of 10 L/min. Acquisitions were averaged and the buffer baseline was subtracted with SpectraManager (JASCO). No smoothing was applied. Data are presented as delta epsilon (Δε) per residue (L·mol-1·cm-1·residue-1) calculated using the molar concentration of peptide and number of residues. Secondary structure content was estimated with BeStSel (https://doi.org/10.1093/nar/gky497) and Selcon3 algorithms using SP175 reference dataset (https://doi.org/10.1002/pro.4153).

### Biolayer Interferometry (BLI) assays

BLI experiments were performed in triplicates on an OctetRED384 instrument (Sartorius) at 25°C. NTA sensors were activated with NiCl2 10mM and loaded either with SFPQ or Streptavidin-His6 (Abcam). Loaded sensors were then incubated with LpDot1 at concentrations ranging from 0.76 to 24.5µM. Specific LpDot1 over SFPQ BLI signals were determined by subtracting the non-specific signals measured on the Streptavidin-loaded sensors. Steady-state analysis was performed using the Octet Analysis Studio software 12, allowing to determine dissociation equilibrium constant (Kd) of the SFPQ/LpDot1 complex.

### Western blot analyses

Sample proteins were prepared in Laemmli sample buffer containing 400 mM β-mercaptoethanol and loaded on SDS PAGE gels, followed by a transfer onto a PVDF membrane (0.2 µm; Trans-Blot Turbo system, Biorad). Amido black staining was carried out to ensure equal protein loading. Membranes were blocked with 5% non-fat milk in TBS-Tween 0.5% for 1 hour and incubated with either anti-SFPQ (Invitrogen, PA5-19663) or anti-histone H3 (Cell Signaling 9715S) primary antibodies overnight at 4 °C. Membranes were washed twice and probed with stabilized goat anti-rabbit horseradish peroxidase-coupled antibody against rabbit IgG (Pierce 1858415) for 1 hour. The proteins were visualized by chemiluminescence detection using HRP Substrate spray reagent (Advansta) on the iBright instrument (ThermoFisher). Images were processed and quantified using Fiji software (ImageJ v2.9.0/1.53t). For quantification, signal intensities of SFPQ between 70-100 kDa were measured and normalized to the corresponding histone H3 loading control. All uncropped and unprocessed scans of the western blots shown in the main and supplemental figures are provided in the **Source Data files**.

### Paraspeckle visualization

For all infection assays with human cells, we used RPMI 1640 Medium, no phenol red supplemented with 2% heat-inactivated FBS. Two hundred thousand dTHP-1 or hMDM cells were seeded in a 96 well round glass bottom µ-Plate (Ibidi 89607) and left uninfected or infected (WT or Δ*lpdot1* mutant *L. pneumophila* strains) for 6- and 12 hours. After 1 h of infection, extracellular bacteria were removed by washing three times with PBS. At the indicated time points, cells were fixed with 4 % paraformaldehyde (PFA, EMS 15714). smFISH was performed essentially as described^84^ with minor modifications. Cells were permeabilized in 70% ethanol (SigmaAldrich 51976) overnight at −20°C. Primary probe sets for the core region of NEAT1 were designed using the Oligostan software^84^ and synthesized as 46 pooled oligonucleotides (IDT oPools) carrying barcode sequences (Oligo IDs from 21 to 65, **Table S2**). Hybridization was carried out overnight at 37°C in buffer containing 20% formamide (Ambion AM9342), 2× SSC (Ambion AM9770), 10% dextran sulfate (SigmaAldrich D8906), and 1 mg/ml tRNA (Roche 10109541001). Following washes in 2× SSC/20% formamide, barcodes were detected using readout probes together with complementary imager labelled with 5’-Cy3/3’-Cy3Sp (IDs 66 and 67, respectively, **Table S2**). SFPQ was stained using the primary rabbit polyclonal antibody (Invitrogen, PA5-29948) and a secondary AlexaFluor 647 donkey anti-rabbit IgG (H+L) (ThermoFisher, A31573). Nuclei were counterstained with DAPI (Sigma, 32670) and EMS Glycerol Mounting Medium with DABCO (EMS 17989-50) was used as a mounting medium. Twelve fields/well were imaged using the high-content screening system Opera Phenix Plus (Revvity), in widefield mode with a 20x/NA 1 water immersion objective, for an XY image resolution of 0.296 µm. The following excitation laser/emission filter (in nm) were used for: DAPI: 405/435-480; Cy3: 561/570-630; EGFP: 488/500-550; Alexa: 647 640/650-760. Images were analyzed with the Signals Image Artist (SImA) software (Revvity). Cells were segmented thanks to the DAPI staining, and only whole cells (*i.e.* not touching image borders) were kept for further analysis. Intracellular EGFP-expressing bacteria and NEAT1 spots / aggregates were also segmented. In both infected and non-infected cells, we measured and analyzed the mean value/well of three parameters coming from inside the nucleus: Alexa 647 and Cy3 intensities (reflecting SFPQ and NEAT1 levels, respectively), and the area of NEAT1 spots/aggregates (to evaluate paraspeckle morphology). Five and three-four biological replicates were considered for dTHP1 and hMDM, respectively. In addition of the mean value/well, single-cell values for the hMDM model were extracted to address the diversity of phenotypes at two levels: (*i*) distributions of individual cells, plotted with each donor represented by a distinct symbol, and (*ii*) donor-level statistics, calculated as mean ± SD across all single cells from the same donor. Statistical comparisons across donors were performed using the Wilcoxon–Mann–Whitney post hoc tests (BH-adjusted p-value). Data represent the mean of four biological replicates, with each donor indicated by a distinct symbol. Normality was assessed using the Shapiro–Wilk test, and statistical comparisons were performed using ordinary one-way ANOVA.

### Cell sorting, RNA isolation, real-time reverse transcription-PCR (RT-PCR) and RNAseq analyses

THP-1 cells were left uninfected or infected with either WT or Δ*lpdot1* mutant *L. pneumophila* strains, both expressing GFP. After 4- or 12 h, cells were sorted by FACS (S3e, BIORAD) according to their GFP-positivity in infection and GFP-negativity for uninfected cells (NI condition), as previously described^85^. Four independent experiments were performed. RNA was extracted from 4.10^6^ cells using the Rneasy Mini Kit (Qiagen) according to the manufacturer’s instructions. RNA samples were treated with DNase (Qiagen) for 15 min at 25 °C and precipitated by sodium acetate and ethanol treatment. RNA concentration was determined by both Nanodrop and QuBit and all samples were checked for the absence of genomic DNA contamination by performing a PCR with the housekeeping gene RPLP0 (Primers IDs 7 and 8 of **Table S2**). The integrity of the RNA samples was measured with the Agilent 2100 bioanalyzer (Agilent Technologies). Libraries were built using an Illumina Stranded mRNA library Preparation Kit (Illumina, USA) following the manufacturer’s protocol. Sequencing was performed on a NextSeq2000 (Illumina, USA) to obtain 108 base paired end reads. cDNA was obtained from 1 ug of RNA using SuperScript IV Reverse Transcriptase (ThermoFisher) using Platinum SYBR Green qPCR SuperMix-UDG (Invitrogen). RT-qPCR was performed on Applied Biosystem by life technologies machine and relative expression levels of splicing isoforms were quantified using isoform-specific primers for NFkB2 and NPIPB11 (Primers IDs from 9 to 12 of the **Table SS2)**. Cycle threshold (Ct) values were normalized to the housekeeping gene RPLP0 to obtain ΔCt values. The proportion of each isoform was calculated according to the following formulas: % Isoform 1=2^−ΔCt1^/(2^−ΔCt1^+2^−ΔCt2^) and % isoform 2=2^−ΔCt2^/(2^−ΔCt1^+2^−ΔCt2^). This approach allowed the relative quantification of isoform usage independent of total transcript abundance^86^. Normality was assessed using the Shapiro–Wilk test, and statistical comparisons were performed using ordinary one-way ANOVA.

### Alternative Splicing Analysis

Illumina sequencing reads were mapped to the human reference genome using STAR aligner (version 2.7.10a) with GRCh38 release 46 primary assembly as reference. Mapping was performed using two-pass alignment mode (--twopassMode Basic) to improve novel junction detection and end-to-end alignment strategy (--alignEndsType EndToEnd) to ensure complete read mapping. Alternative splicing events were identified and quantified using the MAJIQ v2 workflow version 2.5.1^87^. First, the MAJIQ build module was executed using GRCh38 release 46 genome annotations to construct splice graphs from the previously mapped reads. Differential splicing analysis between two conditions was performed using MAJIQ deltapsi to quantify changes in percent spliced-in (Ψ) values. Then, results were filtered using Voila tsv with a minimum threshold of 0.1 for differential splicing (delta-Psi) and a probability threshold of 0.95 to ensure high confidence in the detected events. To simplify the complex splicing patterns and associate the local splicing variations (LSV) to characterize alternative splicing events, the Voila modulize tool, from the MAJIQ software package, was used. In accordance with the previous step, parameters set to identify significant changes between groups (--changing-between-group-dpsi 0.1), with high confidence (--probability-changing-threshold 0.95) and appropriate resolution of complex splicing events (--decomplexify-deltapsi-threshold 0.1). Tables with the details about type of splicing event, *de novo* events along with ΔPSI are shown in **Source Data supplementary file.**

### NFkB2 immunostainings

Two hundred thousand human primary monocytes were seeded in a 96-well glass bottom µ-Plate (Ibidi 89607) coated with 2 µg/ml Poly-D lysine (ThermoFisher, A3890401). Cells were left uninfected or infected with GFP-expressing WT or *Δlpdot1* mutant *L. pneumophila* strains. At the indicated timepoint, cells were fixed with 4 % PFA, incubated for 10 minutes at room temperature with human FcR blocking reagent (Miltenyi, 130-059-901) diluted in PBS containing 0.2% saponin and 2% BSA to prevent nonspecific binding. Cells were incubated with a rabbit monoclonal anti-NFκB2 antibody (Sigma, ZRB1250-4). Following a single wash, cells were incubated with AlexaFluor 647-conjugated donkey anti-rabbit IgG (H+L) secondary antibody (ThermoFisher, A31573), DAPI (Sigma, 32670) to visualize nuclei, and TRITC-conjugated phalloidin (R&D, 5783) to label the actin cytoskeleton. Samples were imaged using the high-content screening system Opera Phenix Plus (Revvity) in widefield mode, with a 20×/NA 1 water immersion objective and twelve images per well were acquired with a resolution of 0.296 µm in both X and Y dimensions. The following dyes excitation laser/emission filter (in nm) were used: DAPI: 405/456; Cy3: 561/599; EGFP 488/522; Alexa 647: 640/706. Images were analysed using the Signals Image Artist (SImA) software (Revvity). Morphological features were extracted using modules that defined nuclear and cytoplasmic boundaries based on DAPI and phalloidin-TRITC staining, respectively. Cells touching the image edges were excluded from analysis. To distinguish infected from non-infected cells, the GFP signal was used to identify intracellular bacteria and calculate infection rates. In both infected and non-infected populations, the median fluorescence intensity of AlexaFluor 647 (NFκB2) was measured in the nucleus and cytoplasm. The ratio of nuclear to cytoplasmic fluorescence intensity was calculated to assess NFκB2 nuclear translocation. Normality was assessed using the Shapiro–Wilk test, and statistical comparisons were performed using ordinary one-way ANOVA.

### CD45RO Flow cytometry analysis

Two hundred thousand human primary monocytes were either left uninfected or infected for 5 hours at MOI 10 with GFP-expressing WT or *Δlpdot1* mutant *L. pneumophila* strains. Following infection, cells were washed with PBS and incubated for 10 minutes at room temperature with human FcR blocking reagent (Miltenyi, 130-059-901) diluted in MACS BSA stock solution (Miltenyi, 130-091-376) diluted [1:20] in AutoMACS Rinsing Solution (#130-091-222). Without additional washing, cells were stained with PE anti-human CD45 recombinant antibody (clone QA17A19, BioLegend, 393412) and APC Fire 750 anti-human CD45RO antibody (clone UCHL1, BioLegend, 304250) and cells incubated on ice for 40 minutes. Cells were then washed with Cell Staining Buffer (BioLegend, 420201), fixed with 4% PFA, and resuspended in 400 μl of Cell Staining Buffer. Samples were acquired on an ID7000 spectral cell analyzer (Sony). Cells were gated based on morphology (SSC-A vs. FSC-A), single-cell discrimination (SSC-A vs. SSC-W), and stratified into GFP-positive (infected) and GFP-negative (uninfected) populations (gating strategy in **Fig. S8**). For both populations, total CD45 (PE) and CD45RO (APC Fire 750) fluorescence intensities were measured. Data were analyzed with FlowJo software, and CD45RO fluorescence was normalized to total CD45 fluorescence using the geometric mean. Normality was assessed using the Shapiro–Wilk test, and statistical comparisons were performed using ordinary one-way ANOVA.

## Notes

### Competing Interest Statement

The authors have declared no competing interest.

