## Supplemental Methods and Figures for "Bacterial targeting of host paraspeckles uncovers a new SFPQ-based regulation"

- **SUPPLEMENTARY FIGURES**

**Figure S1:** Domain alignment of DOT1 homologs

**Figure S2:** Structural characterization of LpDot1

**Figure S3:** Mass spectrometry of LpDot1 targets

**Figure S4:** Modified peptide MS/MS spectrum of SFPQ and its structural characterization in solution

**Figure S5:** Characterization of the LpDot1-SFPQ complex association/dissociation

**Figure S6:** Impact of LpDot1 in virulence and in paraspeckle organization

**Figure S7:** Alternative splicing analyses during *L. pneumophila* infection

**Figure S8:** Gating strategy for the quantification of CD45-PE andCD45RO-APC in infected primary monocytes

**Figure S9:** STRING network analysis of the LpDot1 mediated methylation targets and GO analysis of the differentially spliced genes in WT- and *Δlpdot1* infected cells at 12 h

- **SUPPLEMENTARY MATERIAL AND METHODS**

- **SUPPLEMENTARY TABLES**

**Table S1.** X-ray diffraction data processing and model refinement statistics

### SUPPLEMENTARY FIGURES

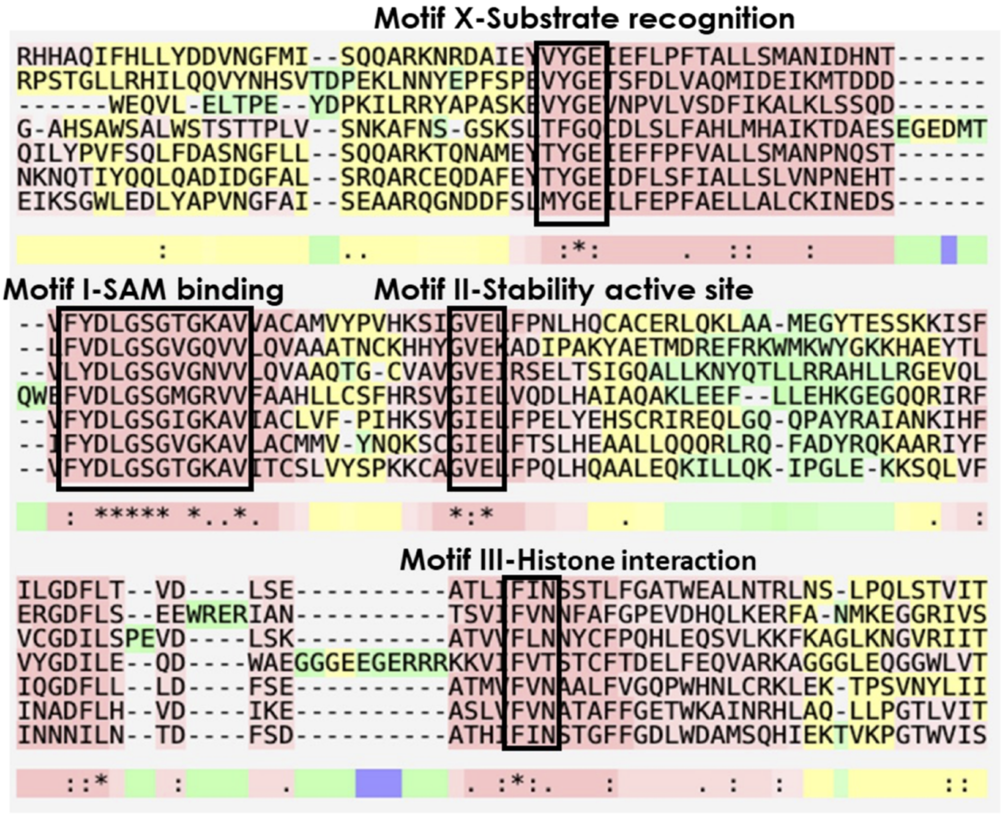


**Figure S1: Domain alignment of DOT1 homologs.** T-Coffee alignment showing the conservation among LpDot1 and the catalytic domains of different eukaryotic proteins. The colour scheme reflects the reliability of the alignment, with residues shaded from red (highest confidence) through yellow and green to blue (lowest confidence). Conservation is indicated according to the Clustal convention, where an asterisk (“*”) denotes fully conserved residues, a colon (“:”) indicates strong conservation among residues with similar physicochemical properties, and a period (“.”) represents weaker conservation. The four black boxes correspond to the conserved motifs of DOT1 methyltransferase domain consisting in: *i*) Motif-X, involved in substrate recognition, *ii*) Motif-I (DxGxGxG), necessary for SAM, *iii*) Motif-II, working synergistically with Motif-I, and *iv*) Motif-III, that plays a role in histone interaction and enzymatic stability.

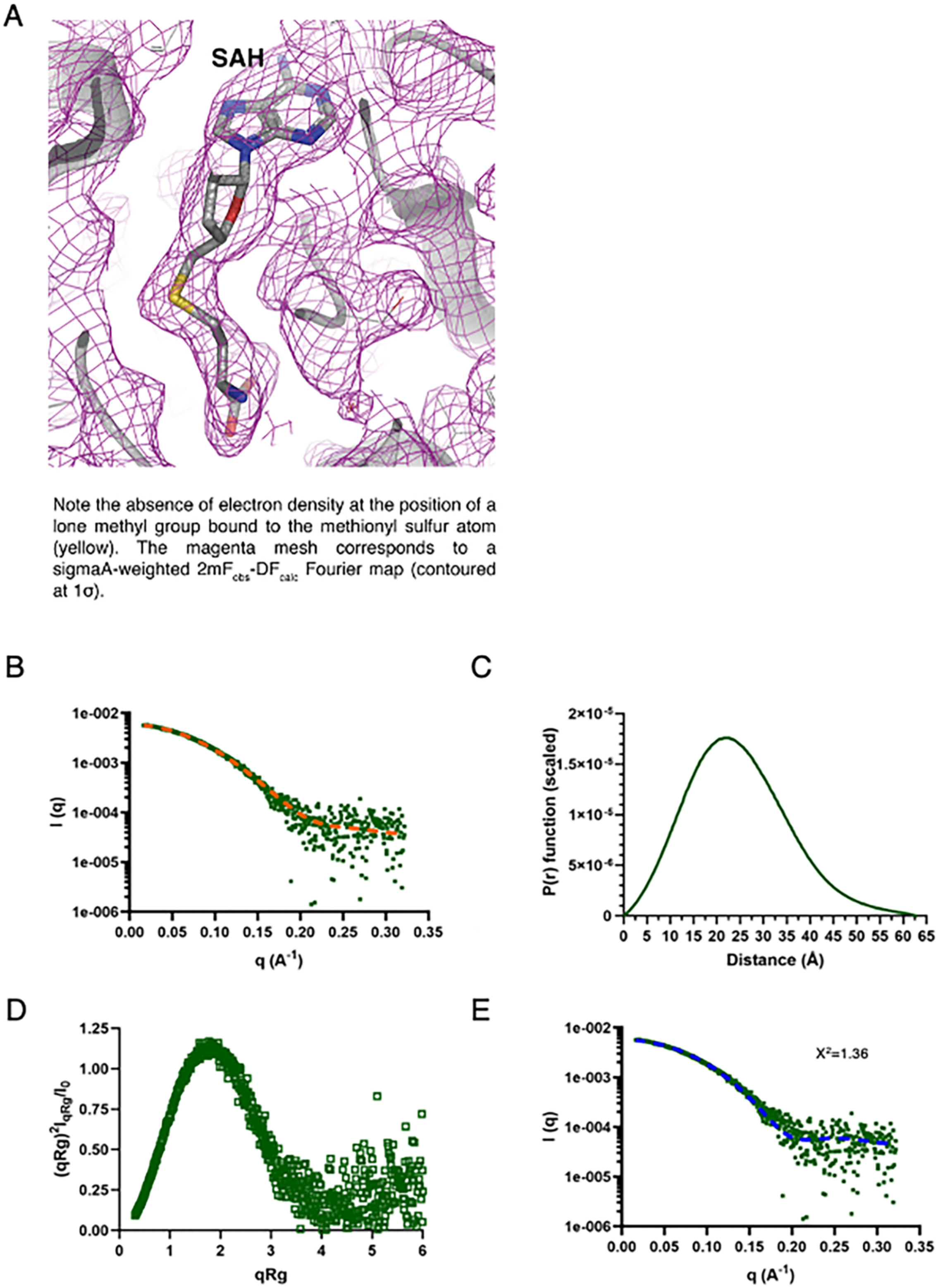


**Figure S2: Structural characterization of LpDot1.** (**A**) Methyl transfer reactions occurred during LpDot1 crystallization. Despite being co-crystallized with SAM, S-adenosyl-L-homocysteine (SAH) was bound within the active site of LpDot1. Note the absence of electron density at the position of a lone methyl group bound to the methionyl sulfur atom (yellow). The magenta mesh corresponds to a sigmaA-weighted 2mFobs- DFcalc Fourier map (contoured at 1σ). **(B-E)** SAXS characterization of LpDot1 in solution. (**B**) Scattering curve of LpDot1(square) with fitted curve (dotted line) used to generate distance distribution. (**C**) Distance distribution functions obtained using the program GNOM. (**D**) Dimensionless Kratky plot. (**E**) Fit of the Crysol calculated scattering pattern using LpDot1 structure.

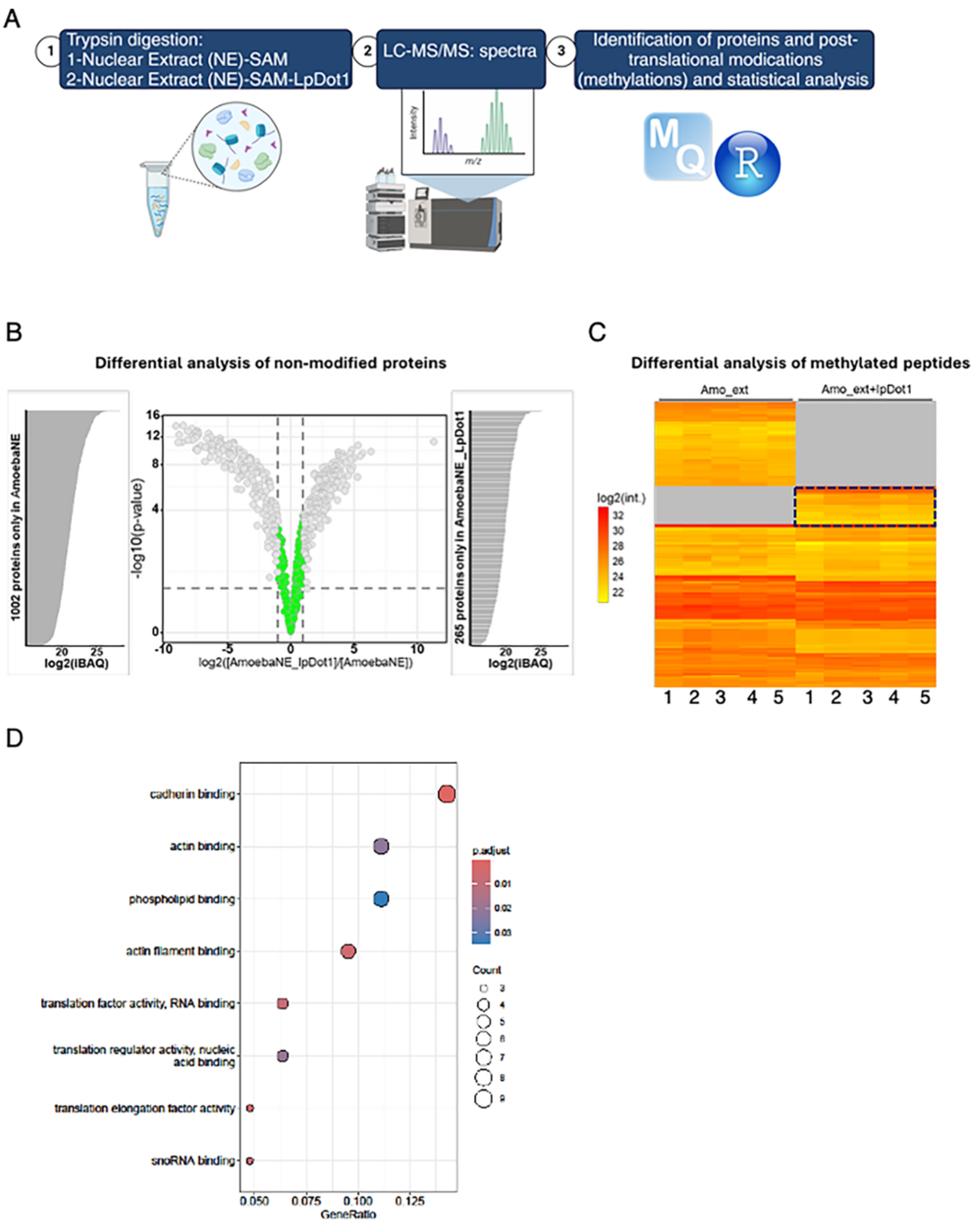


**Figure S3: Mass spectrometry of LpDot1 targets.** (**A**) Workflow depicting **Fig 2B** experiment for the identification of the differentially methylated peptides in NE-SAM compared to NE-SAM-LpDot1 conditions. (**B**) Liquid chromatography and mass spectrometry

(LC-MS) identifying LpDot1 targets in *A. castellanii* nuclear extracts. Volcano plot of -log10 (p values) *vs* log2 values (AmoebaNE-SAM-LpDot1/Amoeba NE-SAM). Green dots represent proteins which abundance is not different between the two groups of comparison. Gray dots are proteins that are differentially abundant in the two conditions and are represented on vertical bar plots on each side. Results from five independent biological replicates. (**C**) Heat map showing the differentially methylated peptides in NE-SAM compared to NE-SAM-LpDot1 (n=5). Black box indicates the peptides showing a methylation that is LpDot1-dependent. (**D**) GO analysis of the biological processes enriched in differentially methylated proteins for which we identified human homologs.

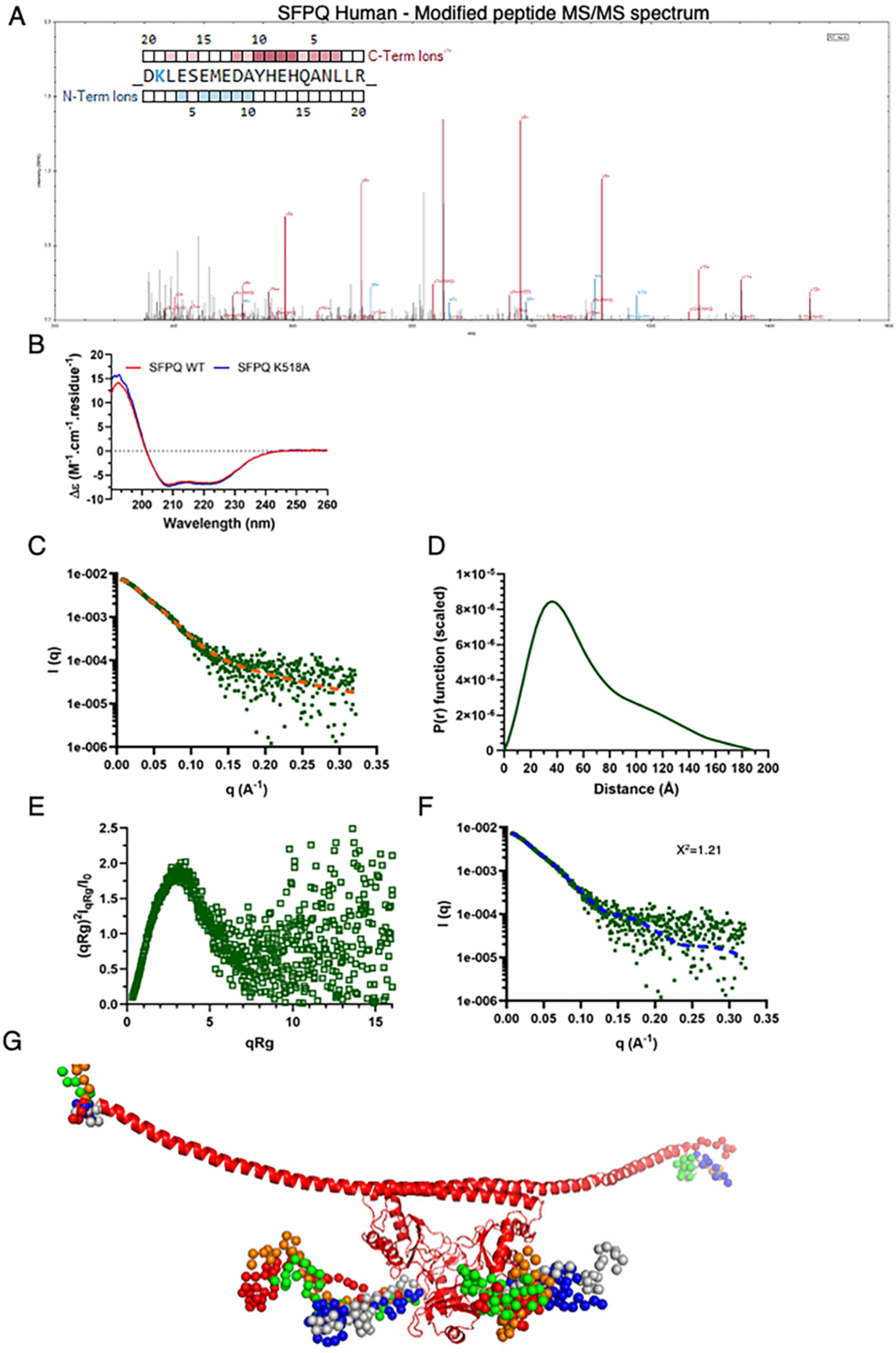


**Figure S4: Modified peptide MS/MS spectrum of SFPQ and its structural characterization in solution.** (A) Fragmentation spectra of SFPQ modified peptide with one PTM: Dimethylation. Highlighted peaks represent peptide fragments. b-ions show fragments from the peptide N-terminus side (blue) and y-ions show fragments from the peptide C-terminus (red). Fragment with modification is highlighted with *: y18. The peptide is dimethylated on lysine. The inset figure at the top-left shows all fragment ions detected. **(B)** Circular dichroism

SFPQ and SFPQ K518A with two negative peaks at ~222 nm and ~208 nm, indicating the prevalence of alpha-helices. **(C-G)** SAXS characterization of SFPQ in solution. **(C)** Scattering curve of SFPQ (square) with fitted curve (dotted line) used to generate distance distribution. **(D)** Distance distribution functions obtained using the program GNOM. **(E)** Dimensionless Kratky plot. **(F)** Fit of the best Coral model. **(G)** Representation of the five best Coral models of SFPQ based on the 4WIJ structure. The structure of the intrinsically disorder regions are represented by spheres.

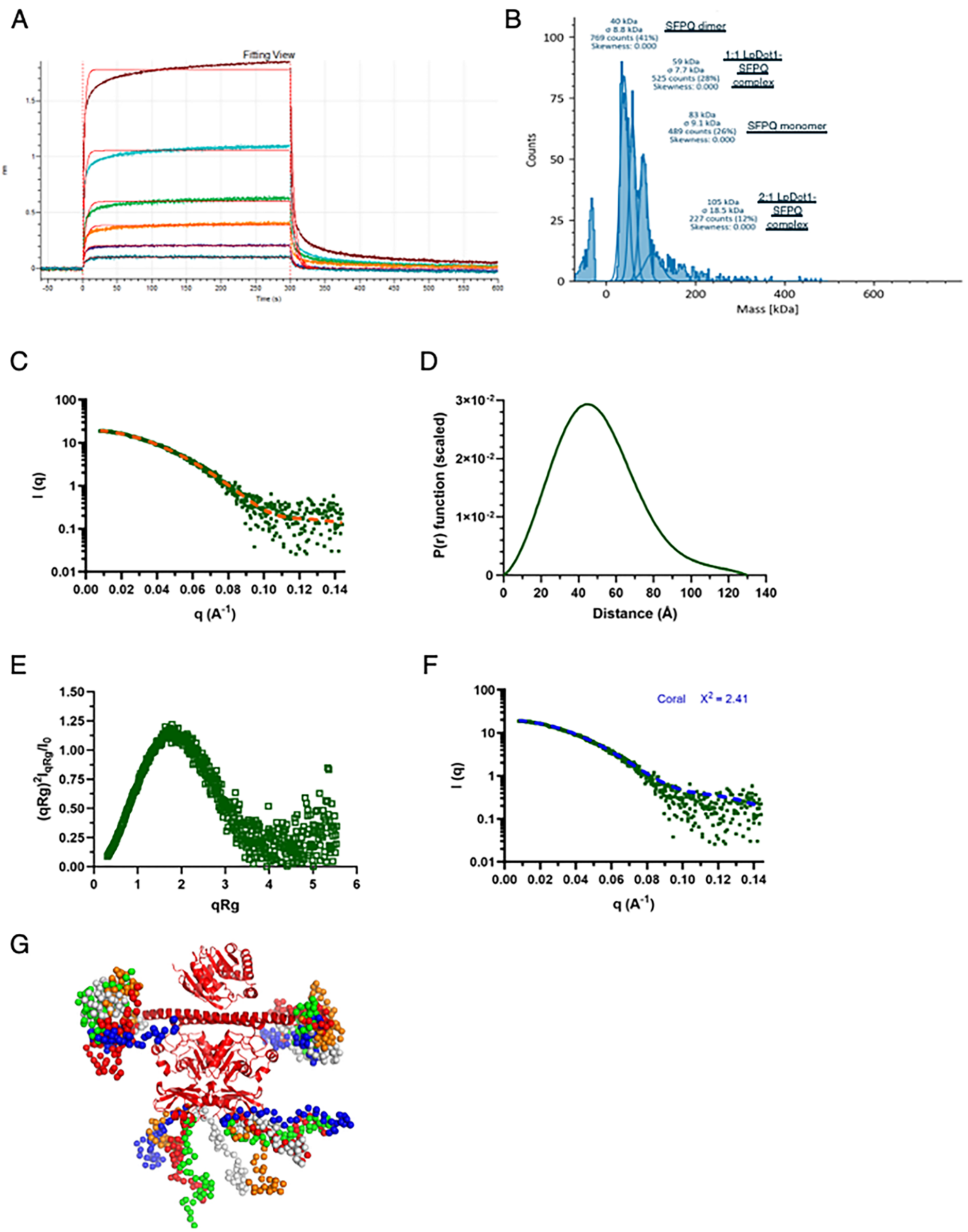


**Figure S5: Characterization of the LpDot1-SFPQ complex association/dissociation. (A)** Representative BLI sensorgram showing the binding interaction between LpDot1 and SFPQ. NTA sensors were loaded either with SFPQ or Streptavidin-His6.Loaded sensors were then incubated with LpDot1 at concentrations spanning from 0.76 to 24.5µM (from gray blue to dark red). Specific LpDot1 over SFPQ BLI signals were determined by subtracting the non-specific ones measured on the Streptavidin-loaded sensors. **(B)** Mass photometry of a 4 nM mixture of LpDot1 with SFPQ (snapshot of the first 2 minutes of interaction). (**C-F**) SAXS characterization of SFPQ-LpDot1 complex in solution. **(C)** Scattering curve of SFPQ-LpDot1 complex (square) with fitted curve (dotted line) used to generate distance distribution. **(D)** Distance distribution functions obtained using the program GNOM. **(E)** Dimensionless Kratky plot. **(F)** Fit of the best Coral model (blue). **(G)** Presentation of the five best Coral models of SFPQ-LpDot1 complex based on the 7UK1. The structure of the intrinsically disorder regions are represented by spheres.

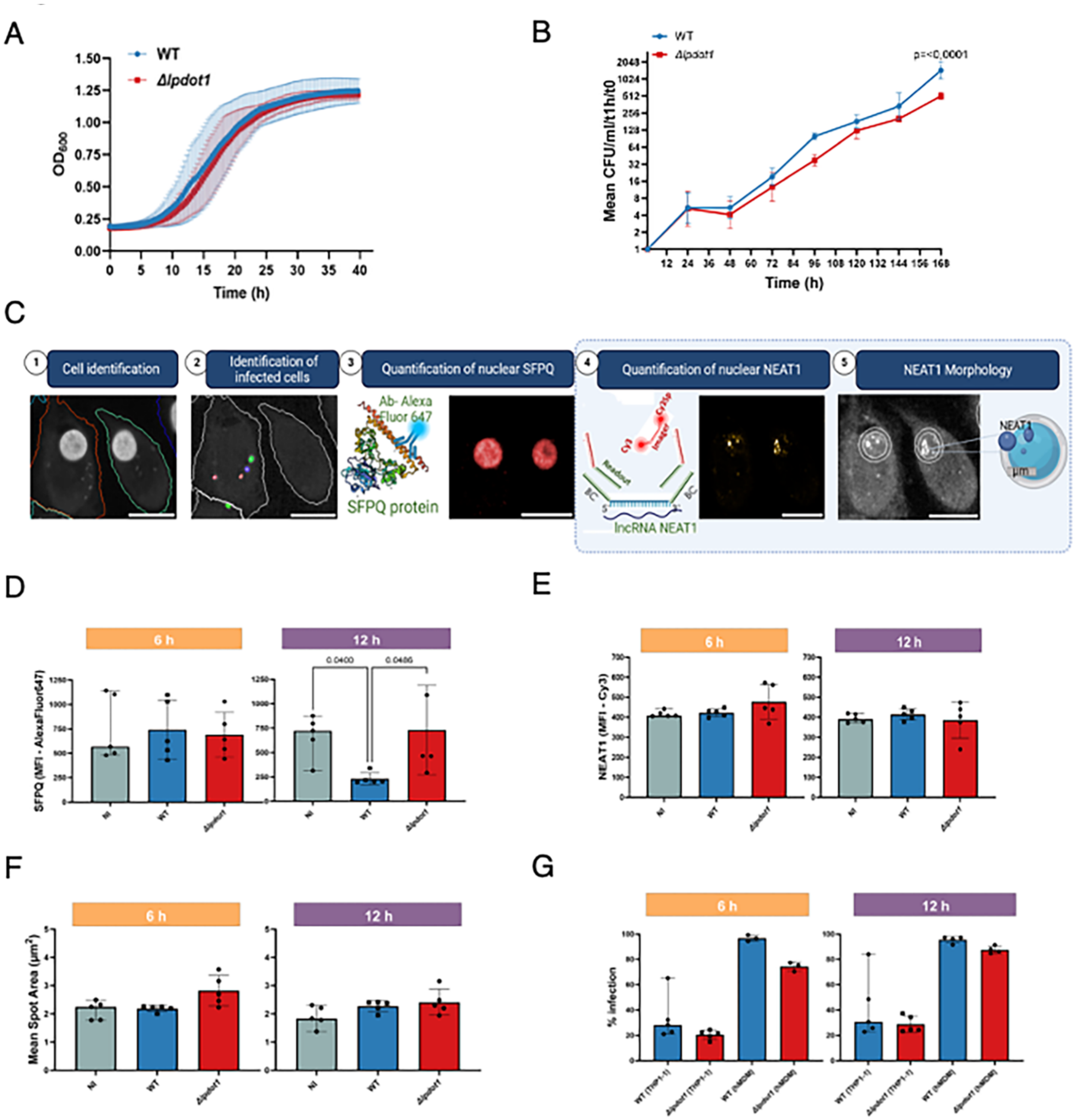


**Figure S6: Impact of *lpdot1* gene in paraspeckle organization.** (**A**) Bacterial growth replication of *L. pneumophila* WT and *Δlpdot1* mutant strains in BYE broth at 37°C. The absorbance at $OD_{600}$ was recorded every 20 min for 39.6 h.(**B**) Intracellular replication of *L. pneumophila* WT and *Δlpdot1* mutant strains in *Acanthamoeba castellanii* (MOI=0.1, 20 °C) was determined by recording the number of colony forming units (CFU) through plating on buffered charcoal yeast extract (BCYE) agar (log2 ratio cfu/t1h/t0). Results are reported as a mean ± SD (n = 3). **(C)** Workflow for the combined RNA-FISH and immunofluorescence analyses used to evaluate paraspeckle content and dynamics. ①-the single cell identification (DAPI signal); ② -selection of infected cells based on the signal from GFP-expressing

*Legionella;* ③- Evaluation of the fluorescence intensity of SFPQ in the nucleus (anti- SFPQ + anti-rabbit Alexa Fluor 647 conjugated); ④- Evaluation of the fluorescence intensity of NEAT1 in the nucleus ( pool of 46 oligo carrying a barcode that is recognized by a reader probe carrying a sequence to which an imager molecule is annealing. The Imager contains a Cy3 molecule that allows NEAT1 RNA to be detected. ⑤- To evaluate the paraspeckle morphology, the area ($\mu m^2$) of NEAT1 positive spots were measured. **(D-F)** Analysis of SFPQ and NEAT1 contents in dTHP-1 cells left uninfected (NI) or infected with either *L. pneumophila* WT or Δ*lpdot1* GFP-expressing strains (MOI=10), for 6- and 12 hours. **(D)** Mean Fluorescence Intensity (MFI) of AlexaFluor 647 (SFPQ content) **(E)** MFI of Cy3 (NEAT1 levels) and **(F)** NEAT1-positive spots area ($\mu m^2$). Results are presented as mean ± SD across 5 independent biological replicates. Statistical comparisons across donors were performed using the Kruskal–Wallis test followed by Dunn's multiple comparison test. **(G)** Evaluation of the percentage on infection at 6- and 12 hours of THP-1 and hMDM cells infected with either *L. pneumophila* WT or Δ*lpdot1* GFP-expressing strains (MOI=10).

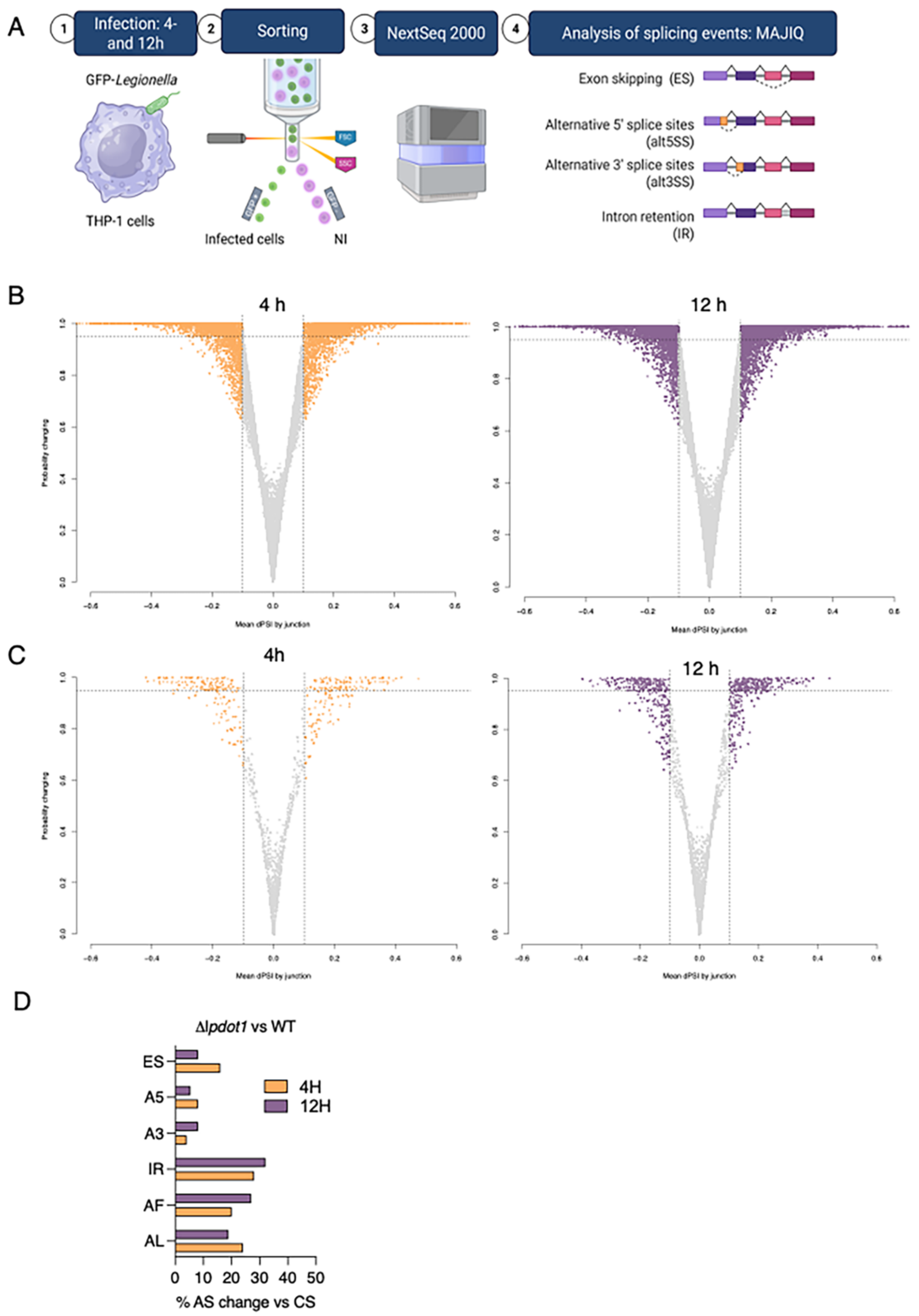


**Figure S7: Alternative splicing analyses during *L. pneumophila* infection. (A)** Workflow depicting the analysis of splicing events in THP-1 cells. Cells were uninfected or infected with either *L. pneumophila* WT or *Δlpdot1* strains expressing GFP. Cells were sorted based on GFP signal in infection or GFP negativity in case of NI cells. RNA was extracted from these samples and their quality checked. Sequencing was performed on a NextSeq2000 (Illumina) to obtain 108 base pair-end read. Splicing analyses were performed using MAJIQ software. **(B-C)** *L.*

*pneumophila*-induced alternative splicing events. Volcano plots showing alternative splicing events of THP-1 cells at 4 and 12 hpi. (**B**) Cells infected with *L. pneumophila* WT compared to non-infected cells. (**C**) Cells infected with *L. pneumophila Δlpdot1* strain compared to cells infected with *L. pneumophila* WT. N=5 biological replicates. Vertical lines indicate a mean dPSI of 0.1, and the horizontal lines indicate a probability of changing of 0.95. **(D)** Percentage of change of significant alternative splicing events per each type of event between WT- and *Δlpdot1*- infected THP1 cells at 4 and 12 hpi.

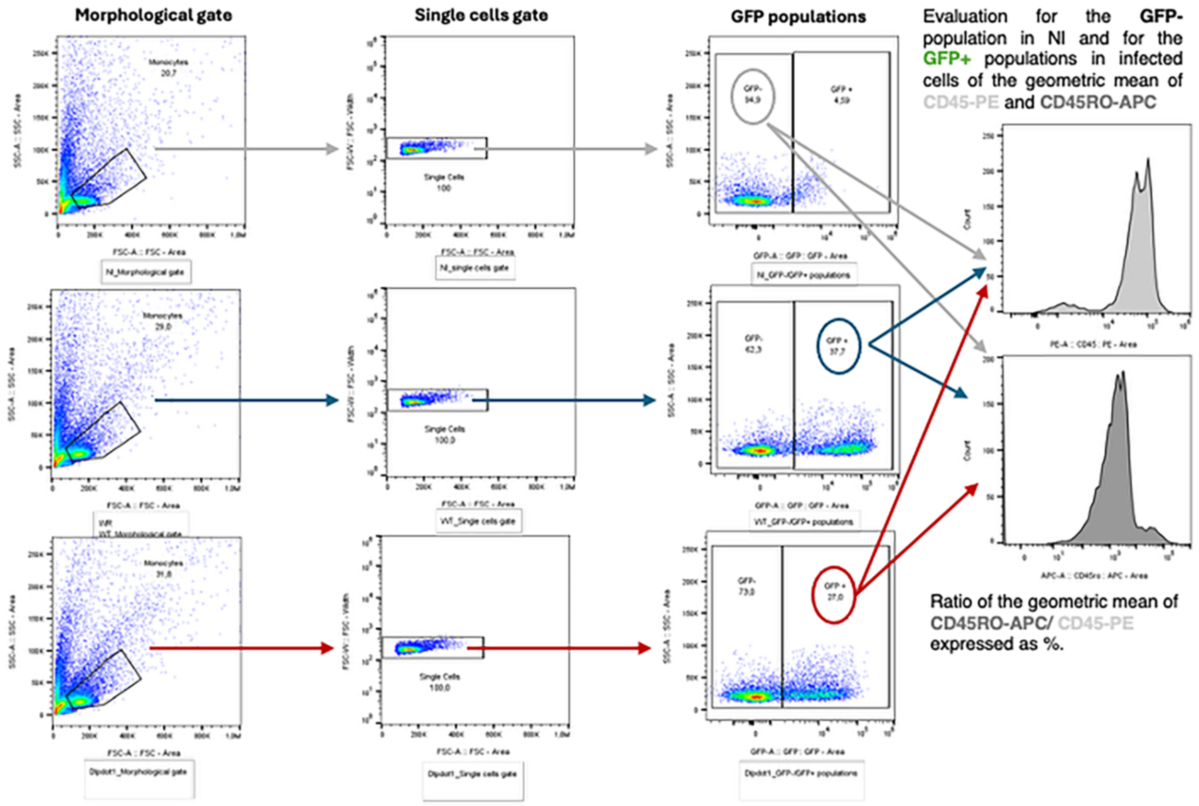


**Figure S8: Gating strategy for the quantification of CD45-PE andCD45RO-APC in infected primary monocytes.** Primary monocytes were uninfected or infected with either *L. pneumophila* WT or *Δlpdot1* strains expressing GFP. Samples were acquired on an ID7000 spectral analyzer (Sony), and a representative example is shown. Cells were gated based on morphology (SSC-A vs. FSC-A), single-cell discrimination (SSC-A vs. SSC-W), and stratified into GFP-positive (infected) and GFP-negative (uninfected) populations.

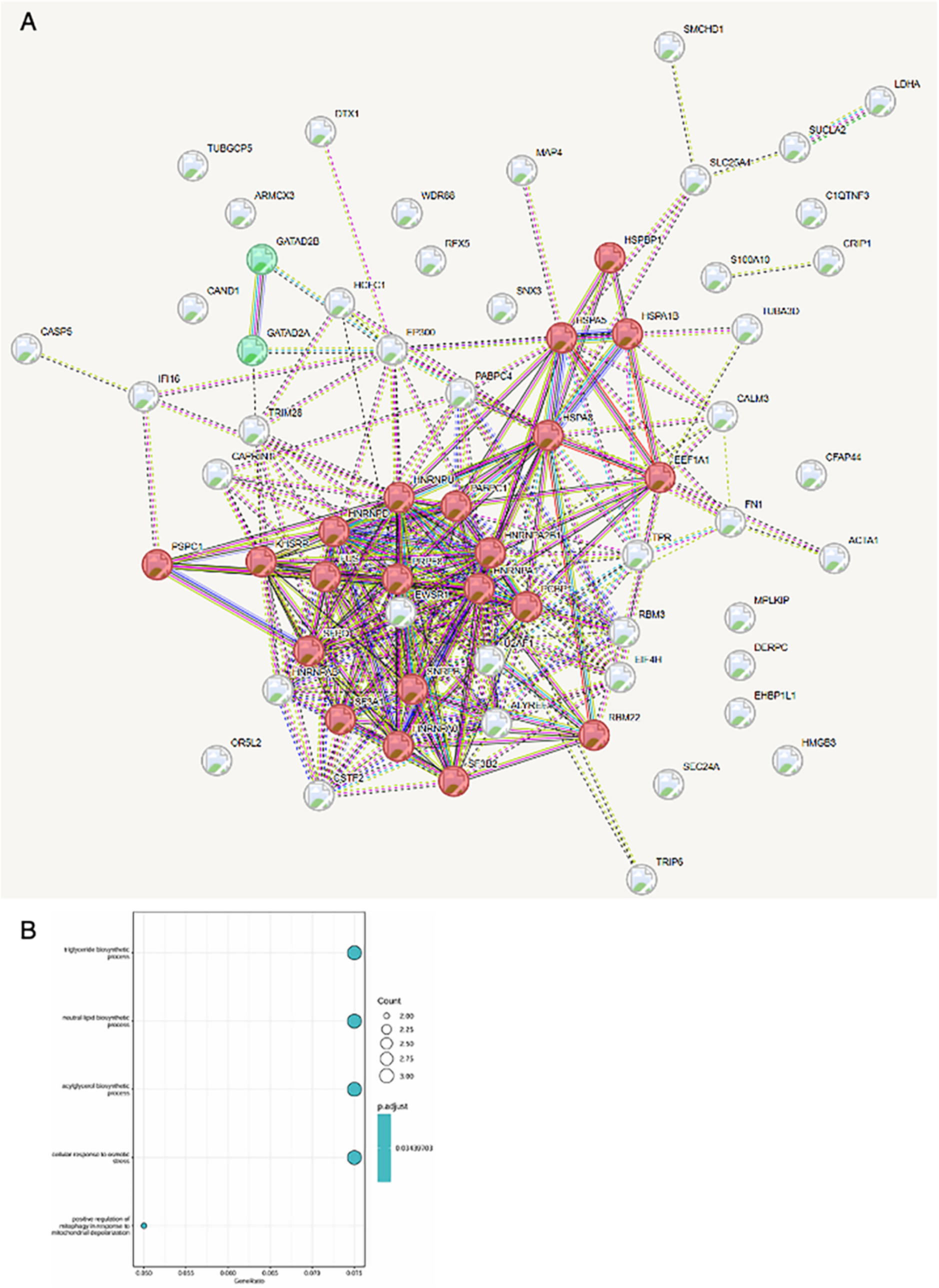


**Figure S9: STRING network analysis of the LpDot1 mediated methylation targets and GO analysis of the differentially spliced genes in WT- and *Δlpdot1* infected cells at 12 h.** (**A**) STRING network analysis of the 64 methylated proteins, based on the **Table S2. (B)** GO analysis of the comparison *Δlpdot1* and WT-infected cells at 12 h post-infection, showing the 5 most enriched biological processes of the genes showing differences in alternative splicing.

### SUPPLEMENTARY MATERIAL AND METHODS

**Protein digestion for human nuclear extract for Tandem Mass Spectrometry (MS/MS) analyses.** Human NE (20 µg) was either incubated or not with 3.5 µg of LpDot1 for 1 h at 30 °C in the MTase reaction buffer (the radioactive SAM was replaced by cold SAM, NEB). For each of the two samples, five biological replicates were prepared. Sample were diluted in 8 M urea, 100 mM Tris HCl pH 8.5 and were reduced using 5mM TCEP for 30 min at room temperature. The alkylation of the reduced disulfide bridges was performed using 10 mM iodoacetamide for 30 min at room temperature in the dark. Proteins were then digested in two steps, first with 1 µg r-LysC Mass Spec Grade (Promega) for 4 h at 30 °C and then samples were diluted below 2 M urea with 100 mM Tris HCl pH 8.5 and 1 µg Sequencing Grade Modified Trypsin was added for the second digestion overnight at 37°C. Proteolysis was stopped by adding formic acid (FA) at a final concentration of 5%. After digestion, samples were acidified with formic acid (FA) and subsequently desalted using an AssayMap C18 cartridges mounted on an Agilent AssayMap BRAVO Liquid Handling System (Agilent). Briefly, C18 cartridges were conditioned first with 100% acetonitrile (ACN) and then with 0.1% FA. Samples were then loaded onto conditioned C18 columns, washed with 0.1% FA, and eluted with 60% ACN, 0.1% FA. Finally, organic solvents were removed in a SpeedVac concentrator prior to LC-MS/MS analysis.

**Liquid chromatography and mass spectrometry (LS-MS) assay**: Dried samples were reconstituted with 2% ACN-0.1% FA and quantified by NanoDrop spectrophometer (Thermo Fisher Scientific). LC-MS/MS analysis was performed on a Q Exactive™ Plus Mass Spectrometer (Thermo Fisher Scientific) coupled with a Proxeon EASY-nLC 1200 (Thermo Fisher Scientific). One µg of peptides was injected onto a home-made 32 cm $C_{18}$ column (1.9 µm particles, 100 Å pore size, ReproSil-Pur Basic C18, Dr. Maisch GmbH, Ammerbuch-Entringen, Germany). Column equilibration and peptide loading were done at 900 bars in buffer A (0.1 % FA). Peptides were separated with a multi-step gradient from 3 to 6 % buffer B (80% ACN, 0.1% FA) in 5 min, 6 to 31 % buffer B in 130 min, 31 to 62 % buffer B in 30 min at a flow rate of 250 nL/min. Column temperature was set to 60°C. MS data were acquired using Xcalibur software using a data-dependent method. MS scans were acquired at a resolution of 70,000 and MS/MS scans (fixed first mass 100 m/z) at a resolution of 17,500. The AGC target and maximum injection time for the survey scans and the MS/MS scans were set to $3E^6$, 20ms and $1E^6$, 60ms respectively. An automatic selection of the 10 most intense precursor ions was activated (Top 10) with a 30 s dynamic exclusion. The isolation window was set to 1.6 m/z and normalized collision energy fixed to 27 for HCD fragmentation. We used an underfill ratio of 1.0 % corresponding to an intensity threshold of $1.7E^5$. Unassigned precursor ion charge states as well as 1, 7, 8 and >8 charged states were rejected and peptide match was disable.

**Protein identification**: Raw data were analyzed using MaxQuant software version v2.0.3.0 [1] with the Andromeda search engine [2] [3]. The MS/MS spectra were searched against the DOT1L_Tag.fasta and the Homo Sapiens Uniprot reference proteome database (79,057 entries). All searches were performed with oxidation of methionine, mono-, di- and trimethylation of lysine, and Acetyl protein N-terminal as variable modifications and cysteine carbamidomethylation as fixed modification. Trypsin was selected as protease allowing for up to two missed cleavages. The minimum peptide length was set to 5 amino acids, and the peptide mass was limited to a maximum of 8,000 Da. One unique peptide to the protein group was required for the protein identification. The main search peptide tolerance was set to 4.5 ppm and to 20 ppm for the MS/MS match tolerance. Second peptides was enabled to identify co-fragmentation events and match between runs option was selected with a match time window of 0.7 min over an alignment time window of 20 min. The false discovery rate (FDR) for peptide and protein identification was set to 0.01. The mass spectrometry proteomics data have been deposited to the ProteomeXchange.

**Statistical analysis of proteomics data**: Statistical analysis compared protein intensities across conditions. Proteins identified as reverse hits or potential contaminants were first removed from the analysis. Intensities associated with at least one unique peptide were kept for further statistics. Additionally, only proteins with at least two intensity values in one of both conditions were kept ensuring a minimum of replicability. Next, intensities were log2-transformed. They were normalized so that the intensities of proteins are centered in each sample on the mean of the medians of the protein intensities across all samples of the same condition. Proteins without any intensity value in one of both conditions have been considered as proteins quantitatively present in one condition and absent in the other. They have, therefore, been set aside and considered as differentially abundant proteins. Next, missing values of the remaining proteins were imputed using the impute.mle function of the R package imp4p [4]. Statistical testing was conducted using limma t-tests thanks to the R package limma [5]. The FDR control was performed using an adaptive Benjamini–Hochberg procedure on the resulting p-values thanks to the function adjust.p of R package cp4p using the robust method to estimate the proportion of true null hypotheses among the set of statistical tests [6]. The proteins associated to an absolute log2(fold-change) superior to 1 and adjusted p-values inferior to a FDR level of 1% have been considered as significantly differentially abundant proteins.

**Statistical analysis of modified peptides and GO analyses**. Post-translational modifications (PTMs) were analyzed. Some modifications were identified by MS1 mass differences, others by MS/MS with localization probabilities >0.75 (MaxQuant) (supplemental material xxx). Raw peptide intensities were log2-transformed and normalized by median centering within biological conditions. Missing values were imputed using impute.mle (R package imp4p). T-tests assessed fold-change significance, and ANOVA contrast analyses were used to highlight modified peptides with fold-changes that deviated significantly from those of their corresponding unmodified proteins [7]. GO term analysis performed by using the GOProfiler in R.

**SFPQ: Samples preparation for Mass Spectrometry (MS).** The full length SFPQ recombinant protein (0.5 µg, Origene) was incubated with or without 1 µg of LpDot1 for 1 at 30 °C in the MTase reaction buffer (the radioactive SAM was replaced by cold SAM, NEB). Samples were digested using S-Trap micro protocol. Samples were diluted with 10% SDS, 50mM TEAB (pH8.5), vortexed, reduced with 5 mM TCEP (15 min at 55°C), and alkylated with 20 mM iodoacetamide (30 min, room temperature, dark). Proteins were precipitated using 2.5% phosphoric acid and six volumes of binding buffer (90% methanol, 100 mM TEAB, pH 7.55). The solution was loaded onto an S-Trap micro filter, centrifuged at 4,000 rpm for 30 s, washed four times, and digested overnight (37°C) with trypsin (1:10 enzyme:protein ratio). Peptides were eluted using stepwise buffers (50 mM TEAB, 0.2% FA in water, and 50% ACN/0.2% FA), dried, and resuspended in 0.1% FA, 2% ACN for LC-MS analysis.

**Liquid Chromatography–MS.** DDA acquisition LC–MS/MS analysis was performed using a nanochromatographic system (Proxeon EASY-nLC 1200 - Thermo Fisher Scientific) coupled online to a Q Exactive Plus Mass Spectrometer (Thermo Fisher Scientific). For each sample, peptides were loaded onto a 50 cm column (EASY-Spray column, 50 cm × 75 µm ID, PepMap C18, 2 µm particles, 100 Å pore size - Thermo Fisher Scientific) after an equilibration step in 100% solvent A ($H_2O$, 0.1% FA). Peptides were eluted using a multi-step gradient: 3 to 6% solvent B (80% ACN, 0.1% FA) over 3 min, 6 to 31% over 82 min, 31 to 62% over 20 min, 62 to 100% over 5 min, and finally from 100% to 3% over 10 min, at a flow rate of 250 nL/min over a total runtime of 120 min. The column temperature was set to 50°C. MS data were acquired using Xcalibur software with a data-dependent Top 10 method. Survey scans (300–1700 m/z) were acquired at a resolution of 70,000, while MS/MS scans (fixed first mass 100 m/z) were acquired at a resolution of 17,500. The AGC target and maximum injection time were set to 3E6 and 20 ms for survey scans and 1E6 and 60 ms for MS/MS scans, respectively. The isolation window was set to 1.6 m/z, and the normalized collision energy was fixed at 28 for HCD fragmentation. A minimum AGC target of 1E4 was used with an intensity threshold of 1.7E5. Unassigned precursor ion charge states, as well as charge states of 1, 7, 8, and >8, were rejected. Peptide match was disabled. The "Exclude isotopes" option was enabled, and selected ions were dynamically excluded for 45 seconds.

**DIA acquisition:** Peptides were injected onto a Thermo Fisher nanochromatographic system (Proxeon EASY-nLC 1200 - Thermo Fisher Scientific) coupled online to an Orbitrap Lumos Mass Spectrometer (Thermo Fisher Scientific). For each sample, peptides were loaded onto a 50 cm column (EASY-Spray column, 50 cm × 75 µm ID, PepMap C18, 2 µm particles, 100 Å pore size - Thermo Fisher Scientific) after an equilibration step in 100% solvent A ($H_2O$, 0.1% FA). Peptides were eluted using a multi-step gradient: 2 to 7% solvent B (80% ACN, 0.1% FA) over 5 min, 7 to 23% over 40 min, 23 to 45% over 25 min, and 45 to 95% over 12 min, at a flow rate of 300 nL/min over a total runtime of 82 min. The column temperature was set to 50°C. The mass spectrometer was operated in DIA mode. A full-scan MS spectrum (m/z 350–1500) was acquired in the Orbitrap with a resolution of 120,000 and a normalized AGC target of 50%.

**MS data processing and analysis.** DDA raw data were analyzed using MaxQuant software version 2.1.4.0 [2,3] using the Andromeda search engine[1]. The MS/MS spectra were searched against the sequence of SFPQ with lpDot1 and usual known mass spectrometry contaminants and reversed sequences of all entries. Andromeda searches were performed choosing trypsin as specific enzyme with a maximum number of two missed cleavages. Possible modifications included carbamidomethylation (Cys, fixed), oxidation (Met, variable), N-terminal acetylation (variable), methyl (Lys/Arg, variable), dimethyl (Lys/Arg, variable), Trimethyl (Lys, variable), Acetyl (Lys, variable) and Acetyl (protein N-term, variable). The mass tolerance in MS was set to 20 ppm for the first search then 4.5 ppm for the main search and 20 ppm for the MS/MS. Maximum peptide charge was set to seven and eight amino acids were required as minimum peptide length. One unique peptide to the protein group was required for the protein identification. A false discovery rate (FDR) cutoff of 1 % was applied at the peptide and protein levels. Quantification was conducted using the XIC-based LFQ algorithm in Fast LFQ mode, as described in [3]. For quantification, unique and razor peptides, including modified peptides, with a minimum of two ratio counts, were accepted.  DIA raw data were analyzed using PEAKS X+ software (PeakS)and Spectronaut (Biognosys). The MS/MS spectra were searched against the sequence of SFPQ with lpDot1. The searches were performed choosing trypsin as specific enzyme. Possible modifications included carbamidomethylation (Cys, fixed), oxidation (Met, variable), N-terminal acetylation (variable), methyl (Lys/Arg, variable), dimethyl (Lys/Arg, variable), Trimethyl (Lys, variable). For PEAKS, the parent mass error tolerance was set to 10 ppm, and the fragment mass error was set to 0.02 DA.

**SUPPLEMENTARY TABLES**

**Table S1.** X-ray diffraction data processing and model refinement statistics.

| | |
|---|---|
| **Data collection** | |
| Space group | P $3_1$ 2 1 |
| *a, b, c* (Å) | a=b=69.44 Å c= 80.77 Å |
| *α, β, γ* (º) | α=β=90º γ=120º |
| Resolution range (Å) | 28.18 – 2.1 (2.22 – 2.11) |
| Nº of unique reflections | 12928 (1803) |
| Completeness (%) | 96.3 (94.2) |
| Multiplicity | 2.9 (2.9) |
| $CC_{1/2}$ | 0.991 (0.16) |
| $R_{meas}$ (%) | 40.3 (294.3) |
| Mean (I / $σ_{(I)}$) | 3.1 (0.4) |
| **Refinement** | |
| $R_{work}$ / $R_{free}$ | 0.217 / 0.270 |
| Number of unique reflections refined against | 11969 |
| Free R value test set count | 992 |
| Nº atoms | |
| Protein | 1613 |
| Ligand (SAH S-adenosyl-homocysteine) | 26 |
| Water | 44 |

| | |
|---|---|
| Deviations from ideal values | |
| Bond lengths (Å) | 0.004 |
| Bond angles (°) | 0.595 |
| Overall B factor from Wilson plot (Å$^2$) | 40.6 |
| All-atom clashscore | 3 |
| Ramachandran plot (% residues) | |
| Outliers | 0 |
| Allowed | 4 |
| Favored | 96 |
| PDB ID | pdb_00009NE1 |

### Table S2. List of primers used in this study

| Primer ID | Primer Name | Sequence (5'-3') | Purpose |
|---|---|---|---|
| 1 | LpDot1_For | GGAAGTTCTGTTCCAGGGGCCCGAACTGCGCATTAATCGCTG | Insertion of LpDot1 (sequence optimized) in pET14 |
| 2 | LpDot1_Rev | TCGAATTCGGATCCGGTACCTCATTATTAGTTCTCTTTTTTATGAATAAAG | Insertion of LpDot1 (sequence optimized) in pET15 |
| 3 | LpDot1 M*For | GTGTTTTACGACCTGCGTAGCCGTACCGGTAAAGCCGTG | Generation of LpDot1 catalytically inactive |
| 4 | LpDot1 M*Rev | CACGGCTTTACCGGTACGGCTACGCAGGTCGTAAAACAC | Generation of LpDot1 catalytically inactive |
| 5 | SFPQK518AFor | catgaaagatgcaaaagacgccttggaaagtgaaatggaagatgcc | Single point mutation for SFPQK518A |
| 6 | SFPQK518ARev | ggcatcttccatttcactttccaaggcgtcttttgcatctttcatg | Single point mutation for SFPQK518A |
| 7 | RPLP0_For | TGGCAGCATCTACAACCCTG | Housekeeping gene qPCR |
| 8 | RPLP0_Rev | AAGGTGTAATCCGTCTCCACAGA | Housekeeping gene qPCR |
| 9 | IntrNFKB2F | GCTCCCCAACCCCCAGAC | NFkB2 qPCR splicing isoform2 |
| 10 | NFKB2F_Ex15R | TGGTTGGTGAGGTTGACA | NFkB2 qPCR splicing isoform1 and isoform 2 |
| 11 | NFKB2F_Ex14F | CGAGCCCTACTCGACTAC | NFkB2 qPCR qPCR splcing isoform2 |
| 12 | NPIPB11_exon2F | CAGCACGTAAAGTCAGTGAC | NPIPB11 qPCR splcing isoform1 |
| 13 | NPIPB11_int2F | CTTGGGAGGTGATGACACTC | NPIPB11 qPCR splicing isoform 2 |
| 14 | NPIPB11_int2R | GTAGCCTACCGTCCATCAAC | NPIPB11 qPCR splcing isoform1 and isoform 2 |
| 15 | Lpp3025upstreamFor | gaagattctctgactatg | Deletion of *lpdot1* in L. *pneumophila* |
| 16 | Lpp3025upstreamRev | gactctagaggatccTTAcatttttgacgatatgaa | Deletion of *lpdot1* in L. *pneumophila* |
| 17 | Lpp3025downstreamFor | AaagcgaatcCCTAGGcataagaaagaaaactaatttttac | Deletion of *lpdot1* in L. *pneumophila* |
| 18 | Lpp3025downstreamRev | Gatgttcagtctgatgtctg | Deletion of *lpdot1* in L. *pneumophila* |
| 19 | MycLpDot1_for | CAGAATTCAGAAGGAGATATACATATGGAACAAAAACTCATCTCAGAAGAGGATCTGATGCTTTTTTTATGCCTCATTC | Complementation of *lpdot1* |
| 20 | MycLpDot1_rev | CCGCCAAAACAGCCAAGCTTAGTTTTCTTTCTTATGAATAAATGC | Complementation of *lpdot1* |
| 21 | h_neat1-middle | AAATTGCGTGACGGACCTGGTCCTAGCATGGCACATGCATATCCTAAATTGCGTGACGGACCTGG | Oligo of the pool of 46 targeting NEAT1 core region and containing an appended sequence that anneals to the readout probe. |
| 22 | h_neat1-middle | AAATTGCGTGACGGACCTGGACACGAGGCACAGTCCTGCATGCTCAAATTGCGTGACGGACCTGG | Oligo of the pool of 46 targeting NEAT1 core region and containing an appended sequence that anneals to the readout probe. |
| 23 | h_neat1-middle | AAATTGCGTGACGGACCTGGTAGTAGGGTGGGATAGGTGAGGGCAGGGATAAATTGCGTGACGGACCTGG | Oligo of the pool of 46 targeting NEAT1 core region and containing an appended sequence that anneals to the readout probe. |
| 24 | h_neat1-middle | AAATTGCGTGACGGACCTGGTCCTGGGAACTGCCATCACTCGAAATGAAATTGCGTGACGGACCTGG | Oligo of the pool of 46 targeting NEAT1 core region and containing an appended sequence that anneals to the readout probe. |

| | | | |
|---|---|---|---|
| 25 | h_neat1-middle | AAATTGCGTGACGGACCTGGTCTGCAGGCCTCAGACCTCTCAAAGAAATTGCGTGACGGACCTGG | Oligo of the pool of 46 targeting NEAT1 core region and containing an appended sequence that anneals to the readout probe. |
| 26 | h_neat1-middle | AAATTGCGTGACGGACCTGGGGCCTTACAAGGCCTCAGAAATGGGGGAAATTGCGTGACGGACCTGG | Oligo of the pool of 46 targeting NEAT1 core region and containing an appended sequence that anneals to the readout probe. |
| 27 | h_neat1-middle | AAATTGCGTGACGGACCTGGGATGTGGAATGTGACAGAGAATCCCTTCCCAAATTGCGTGACGGACCTGG | Oligo of the pool of 46 targeting NEAT1 core region and containing an appended sequence that anneals to the readout probe. |
| 28 | h_neat1-middle | AAATTGCGTGACGGACCTGGAGCAGCAGCGTGAAGGCGAGGCTGTCAAATTGCGTGACGGACCTGG | Oligo of the pool of 46 targeting NEAT1 core region and containing an appended sequence that anneals to the readout probe. |
| 29 | h_neat1-middle | AAATTGCGTGACGGACCTGGCTGAGGGCATGGTGGGGCTAGAGCAAGAAATTGCGTGACGGACCTGG | Oligo of the pool of 46 targeting NEAT1 core region and containing an appended sequence that anneals to the readout probe. |
| 30 | h_neat1-middle | AAATTGCGTGACGGACCTGGGTTGGGAACTTGTCTAGATATTTCCCATCATAAAATTGCGTGACGGACCTGG | Oligo of the pool of 46 targeting NEAT1 core region and containing an appended sequence that anneals to the readout probe. |
| 31 | h_neat1-middle | AAATTGCGTGACGGACCTGGGATCCAGCACATCTAGCAGGGGGTGTAAATTGCGTGACGGACCTGG | Oligo of the pool of 46 targeting NEAT1 core region and containing an appended sequence that anneals to the readout probe. |
| 32 | h_neat1-middle | AAATTGCGTGACGGACCTGGTTGTGCTTCTCATCATTTCACATTAAGAACCCAAATTGCGTGACGGACCTGG | Oligo of the pool of 46 targeting NEAT1 core region and containing an appended sequence that anneals to the readout probe. |
| 33 | h_neat1-middle | AAATTGCGTGACGGACCTGGTTTTGCCATATCTGACTGAATATAGCCAGTCAAATTGCGTGACGGACCTGG | Oligo of the pool of 46 targeting NEAT1 core region and containing an appended sequence that anneals to the readout probe. |
| 34 | h_neat1-middle | AAATTGCGTGACGGACCTGGGCTGTGGACAGCAGGAATAGGCTGTTGAGAAATTGCGTGACGGACCTGG | Oligo of the pool of 46 targeting NEAT1 core region and containing an appended sequence that anneals to the readout probe. |
| 35 | h_neat1-middle | AAATTGCGTGACGGACCTGGACAGGCTAACTAACCCTGTCACTTGTTATGAAATTGCGTGACGGACCTGG | Oligo of the pool of 46 targeting NEAT1 core region and containing an appended sequence that anneals to the readout probe. |
| 36 | h_neat1-middle | AAATTGCGTGACGGACCTGGGGCAGGGCCTTGCTTTGACACACAGAAAATTGCGTGACGGACCTGG | Oligo of the pool of 46 targeting NEAT1 core region and containing an appended sequence that anneals to the readout probe. |
| 37 | h_neat1-middle | AAATTGCGTGACGGACCTGGCCTTCTAGAGGTAGTAAGTCCACCGAGGCAAATTGCGTGACGGACCTGG | Oligo of the pool of 46 targeting NEAT1 core region and containing an appended sequence that anneals to the readout probe. |
| 38 | h_neat1-middle | AAATTGCGTGACGGACCTGGGAGGGCAGGCGCAGACAGCACTCCTAAATTGCGTGACGGACCTGG | Oligo of the pool of 46 targeting NEAT1 core region and containing an appended sequence that anneals to the readout probe. |
| 39 | h_neat1-middle | AAATTGCGTGACGGACCTGGTGCTAAGCCAGGCACCGTGTTATACTCAAATTGCGTGACGGACCTGG | Oligo of the pool of 46 targeting NEAT1 core region and containing an appended sequence that anneals to the readout probe. |
| 40 | h_neat1-middle | AAATTGCGTGACGGACCTGGTAAAGTCCTGGTATAATAGGTGCTTTTTGCACAAATTGCGTGACGGACCTGG | Oligo of the pool of 46 targeting NEAT1 core region and containing an appended sequence that anneals to the readout probe. |

| | | | |
|---|---|---|---|
| 41 | h_neat1-middle | AAATTGCGTGACGGACCTGGTGATTCCCGTTACATGAAAGGTCCAGCGCAAATTGCGTGACGGACCTGG | Oligo of the pool of 46 targeting NEAT1 core region and containing an appended sequence that anneals to the readout probe. |
| 42 | h_neat1-middle | AAATTGCGTGACGGACCTGGGACCTGACGCTAGGCACCAGGGGAGAAATTGCGTGACGGACCTGG | Oligo of the pool of 46 targeting NEAT1 core region and containing an appended sequence that anneals to the readout probe. |
| 43 | h_neat1-middle | AAATTGCGTGACGGACCTGGGGGGAAGGAGGGCAGAGGCAGGAGAGTTCAAATTGCGTGACGGACCTGG | Oligo of the pool of 46 targeting NEAT1 core region and containing an appended sequence that anneals to the readout probe. |
| 44 | h_neat1-middle | AAATTGCGTGACGGACCTGGTTCTGAATTGAACCCTGCCACTGTGACAAAAATTGCGTGACGGACCTGG | Oligo of the pool of 46 targeting NEAT1 core region and containing an appended sequence that anneals to the readout probe. |
| 45 | h_neat1-middle | AAATTGCGTGACGGACCTGGACAGGAGGTGGGTTCTGAGAGCACCATGAAATTGCGTGACGGACCTGG | Oligo of the pool of 46 targeting NEAT1 core region and containing an appended sequence that anneals to the readout probe. |
| 46 | h_neat1-middle | AAATTGCGTGACGGACCTGGGGATGCCTTCACTCAGAGGAAGTTCACAGAAATTGCGTGACGGACCTGG | Oligo of the pool of 46 targeting NEAT1 core region and containing an appended sequence that anneals to the readout probe. |
| 47 | h_neat1-middle | AAATTGCGTGACGGACCTGGCTCAACCAATACTGGCTCTCAGGGTACAAATTGCGTGACGGACCTGG | Oligo of the pool of 46 targeting NEAT1 core region and containing an appended sequence that anneals to the readout probe. |
| 48 | h_neat1-middle | AAATTGCGTGACGGACCTGGCAGCATGCCAGTTGCTGGAGACAGCAAAATTGCGTGACGGACCTGG | Oligo of the pool of 46 targeting NEAT1 core region and containing an appended sequence that anneals to the readout probe. |
| 49 | h_neat1-middle | AAATTGCGTGACGGACCTGGGGATTGTTTCTGGGGCATCCAGGGAAAGGAAAATTGCGTGACGGACCTGG | Oligo of the pool of 46 targeting NEAT1 core region and containing an appended sequence that anneals to the readout probe. |
| 50 | h_neat1-middle | AAATTGCGTGACGGACCTGGGAGTAGAACAAGCCTCCCTTGCTGCTAAATTGCGTGACGGACCTGG | Oligo of the pool of 46 targeting NEAT1 core region and containing an appended sequence that anneals to the readout probe. |
| 51 | h_neat1-middle | AAATTGCGTGACGGACCTGGCTTTGAAACAAGCACCTTCCTTCTCAGCAAAATTGCGTGACGGACCTGG | Oligo of the pool of 46 targeting NEAT1 core region and containing an appended sequence that anneals to the readout probe. |
| 52 | h_neat1-middle | AAATTGCGTGACGGACCTGGCAGTTTGCAAGCCAAGATGTATCCTCCAGAAATTGCGTGACGGACCTGG | Oligo of the pool of 46 targeting NEAT1 core region and containing an appended sequence that anneals to the readout probe. |
| 53 | h_neat1-middle | AAATTGCGTGACGGACCTGGGCACACAGGGAAGCCCAGACGTTTTAGTCAAATTGCGTGACGGACCTGG | Oligo of the pool of 46 targeting NEAT1 core region and containing an appended sequence that anneals to the readout probe. |
| 54 | h_neat1-middle | AAATTGCGTGACGGACCTGGCTATGGAATATGCATATTATGTGTCCATTTCTAAATTGCGTGACGGACCTGG | Oligo of the pool of 46 targeting NEAT1 core region and containing an appended sequence that anneals to the readout probe. |
| 55 | h_neat1-middle | AAATTGCGTGACGGACCTGGGTCTGCAGACCACTTGGGATAATACCAGTAAATTGCGTGACGGACCTGG | Oligo of the pool of 46 targeting NEAT1 core region and containing an appended sequence that anneals to the readout probe. |
| 56 | h_neat1-middle | AAATTGCGTGACGGACCTGGTTTTGCTTTTTCCTGCTCACTCTTTCACAGAAATTGCGTGACGGACCTGG | Oligo of the pool of 46 targeting NEAT1 core region and containing an appended sequence that anneals to the readout probe. |

| | | | |
|---|---|---|---|
| 57 | h_neat1-middle | AAATTGCGTGACGGACCTGGTACTTGCATTTTATACCCATGGCTTGAATTTTAAATTGCGTGACGGACCTGG | Oligo of the pool of 46 targeting NEAT1 core region and containing an appended sequence that anneals to the readout probe. |
| 58 | h_neat1-middle | AAATTGCGTGACGGACCTGGGGACACTTCCGGCACCCTGGCCTAGAAATTGCGTGACGGACCTGG | Oligo of the pool of 46 targeting NEAT1 core region and containing an appended sequence that anneals to the readout probe. |
| 59 | h_neat1-middle | AAATTGCGTGACGGACCTGGGAAATGGTTCTCTGGGTCTGCTGGGCCAAATTGCGTGACGGACCTGG | Oligo of the pool of 46 targeting NEAT1 core region and containing an appended sequence that anneals to the readout probe. |
| 60 | h_neat1-middle | AAATTGCGTGACGGACCTGGTTCTCCTGTCACTTGACTCTAACCTGGAAATTGCGTGACGGACCTGG | Oligo of the pool of 46 targeting NEAT1 core region and containing an appended sequence that anneals to the readout probe. |
| 61 | h_neat1-middle | AAATTGCGTGACGGACCTGGTCCAAGTCACCAGGGCCCCAGGACCAAATTGCGTGACGGACCTGG | Oligo of the pool of 46 targeting NEAT1 core region and containing an appended sequence that anneals to the readout probe. |
| 62 | h_neat1-middle | AAATTGCGTGACGGACCTGGAGGGTGAGAGCTGGGTGCCTTCTACTAAATTGCGTGACGGACCTGG | Oligo of the pool of 46 targeting NEAT1 core region and containing an appended sequence that anneals to the readout probe. |
| 63 | h_neat1-middle | AAATTGCGTGACGGACCTGGGGACTACCAGGTGGGCTCCGAGCCAAAATTGCGTGACGGACCTGG | Oligo of the pool of 46 targeting NEAT1 core region and containing an appended sequence that anneals to the readout probe. |
| 64 | h_neat1-middle | AAATTGCGTGACGGACCTGGGTGGGGAATGTGTTGGGCCAACTGCCAAATTGCGTGACGGACCTGG | Oligo of the pool of 46 targeting NEAT1 core region and containing an appended sequence that anneals to the readout probe. |
| 65 | h_neat1-middle | AAATTGCGTGACGGACCTGGATGGTGCGGGCACTTACTTACTGCAGAAATTGCGTGACGGACCTGG | Oligo of the pool of 46 targeting NEAT1 core region and containing an appended sequence that anneals to the readout probe. |
| 66 | Readout-10 | CCAGGTCCGTCACGCAATTTggccaatggcGATCCGATTGGAACCGTCCCA | Oligo that anneals to the oligo pool and to Imager1 |
| 67 | Imager-1-Cy3-3'-5' | /5Cy3/TG GGA CGG TTC CAA TCG GAT C/3Cy3Sp/ | Oligo that anneals to the Readout-10 labeled with Cy3 molecule |
| 68 | mazF-AprR Forward | CTGTGCTTGATTTCATATCGTCAAAAATGTAAGGATCCACTAGTAACGGCCG | Amplification of the dual marker cassette mazF-AprR |
| 69 | mazF-AprR Reverse | cggccgcgaattccctagggattcgctttccagtcg | Amplification of the dual marker cassette mazF-AprR |
| 70 | Forward primer to check Lpp3025 deletion | Catcaaaggatattattagc | Primer to verify Lpp3025 deletion |
| 71 | Reverse primer to check Lpp3025 deletion | gtatgatgttcgctgttcg | Primer to verify Lpp3025 deletion |
